# Microbiota-derived extracellular vesicles link intestinal dysbiosis to neuroimmune activation in long COVID

**DOI:** 10.64898/2026.02.28.708602

**Authors:** Matheus Aranguren, Kim Doyon-Laliberté, Idia Boncheva, Alexandre Villard, Aléhandra Desjardins, Emma Darbinian, Suhani Patel, Charlotte DuSablon, Estefania Rivera Conde, Diana Cabrera Munoz, Ludhovik Purchase, Valerio E. C. Piscopo, Aeshah Alluli, Faiza Benaliouad, Julien Sirois, Thomas Martin Durcan, Chantal Massé, Kodjovi Dodji Mlaga, Prabha Chandrasekaran, Johanne Poudrier, Emilia Liana Falcone

## Abstract

Post COVID-19 condition (Long COVID, LC) is frequently accompanied by persistent neurological symptoms, but the mechanisms linking intestinal dysbiosis to neuroinflammation remain unclear. Here we identify gut microbiota-derived extracellular vesicles (GMEVs) as functional mediators linking LC-associated dysbiosis to systemic and neuroimmune inflammation. In a longitudinally characterized cohort, individuals with LC and neurological symptoms exhibit a persistent intestinal microbiome signature. Transplantation of LC-associated microbiota into germ-free mice induces intestinal barrier disruption and neuroinflammatory phenotypes. GMEVs from individuals with LC activate inflammasome-associated programs and impair epithelial barrier function, promote inflammatory responses in macrophages, and induce coordinated pro-inflammatory transcriptional programs in human induced pluripotent stem cell (iPSC)-derived microglia. Chronic oral administration of LC-derived GMEVs remodels the microbiota and induces intestinal and systemic inflammation with glial activation *in vivo*. Together, these findings support a vesicle-centered framework in which microbiota-derived extracellular vesicles translate dysbiosis into sustained immune and neuroimmune activation in a post-viral inflammatory state.

## Introduction

Approximately 5-19% of individuals infected with severe acute respiratory syndrome coronavirus 2 (SARS-CoV-2) develop post COVID-19 condition (PCC), also referred to as post-acute sequelae of COVID-19 (PASC) or Long COVID (LC)^1,2^. LC is a post-infectious chronic condition present for at least 3 months after acute SARS-CoV-2 infection, and it may follow a continuous, relapsing-remitting or progressive disease course affecting one or multiple organ systems^2,3^. Although more than 200 symptoms have been attributed to LC^4,5^, some of the most prevalent and debilitating manifestations include fatigue, post-exertional malaise, cognitive impairment (“brain fog”), memory and concentration difficulties, dyspnea and cardiac dysautonomia, including postural orthostatic tachycardia syndrome^6–8^. These sequelae can substantially impair functional capacity and quality of life for years after infection^2,9^. Despite its growing public health burden, LC currently lacks validated diagnostic biomarkers and targeted pharmacological therapies. Accumulating evidence implicates immune dysregulation^10–12^, chronic systemic inflammation^13,14^, microclotting^15^, neuroinflammation^16,17^, autoimmunity^18,19^ and viral persistence^20,21^, including prolonged viral shedding in the gastrointestinal (GI) tract^22–24^, as features of LC pathophysiology.

Acute SARS-CoV-2 infection is associated with intestinal dysbiosis, particularly in individuals with severe disease^25^, and alterations in the gut microbiota can persist for at least 6 months in individuals with LC^26^. However, the mechanisms by which SARS-CoV-2-associated dysbiosis contributes to immune dysregulation and end-organ injury, including neuroinflammation, remain poorly defined. The intestinal microbiota plays a central role in immune homeostasis^27^, particularly at the mucosal barrier^28^, and dysbiosis has been linked to increased intestinal inflammation and epithelial permeability (“leaky gut”), permitting the translocation of microbial products into systemic circulation^29^. Increased intestinal permeability has been documented during acute COVID-19 infection^30,31^, and circulating markers of microbial translocation are elevated in individuals with LC^32^. Although the gut-brain axis has been implicated in neuropsychological and neurodegenerative disorders^33^, including in chronic viral infections such as human immunodeficiency virus (HIV)^34–36^, a causal relationship between intestinal dysbiosis, barrier dysfunction, immune activation, and neuroinflammation has not been demonstrated in LC.

Bacteria communicate with each other and their host through the release of extracellular vesicles (EVs), nanoscale (20-400 nm) lipid bilayer particles that transport proteins, lipids, metabolites and nucleic acids^37^. EVs produced by the intestinal microbiota, here termed gut microbiota-derived extracellular vesicles (GMEVs), can cross the intestinal barrier, enter the bloodstream and act on distant tissues, thereby contributing to diverse inflammatory and metabolic disorders, including metabolic syndrome and liver disease^38–40^. We hypothesized that GMEVs produced by the perturbed microbiota of individuals with LC promote intestinal barrier dysfunction, systemic inflammation and downstream neuroinflammation. Here, we functionally characterized the fecal microbiome of individuals with LC at 3-6 and 12 months post-infection and examined the biological effects of LC-derived GMEVs using complementary *in vitro* and *in vivo* models. Our findings identify GMEVs as functional mediators linking intestinal dysbiosis to barrier dysfunction, immune activation, and neuroinflammatory manifestations in LC.

## Results

### Women with LC and neurological symptoms have intestinal dysbiosis that transfers neurobehavioral alteration and neuroinflammation to gnotobiotic mice

We analyzed a sub-cohort of 103 participants from the *Institut de recherches cliniques de Montréal* (IRCM) Post-COVID-19 (IPCO) cohort (NCT04736732), comprising 12 pandemic controls (PC; individuals who never reported COVID-19 symptoms or tested positive for SARS-CoV-2) and 91 individuals with LC who were not hospitalized during acute infection, all of whom provided stool samples for shotgun metagenomic sequencing (Figure 1A). The median age (interquartile range) of PC and LC participants was 50 (46.5-63) and 47 (39-58) years, respectively, and 41.7% of PC and 64.8% of LC participants self-identified as female. Female sex was associated with higher odds of LC symptom reporting overall (odds ratio (OR) = 1.26, *P* = 6.3 × 10^−9^; Figure 1B) including several neurological symptoms, such as trouble with memory (OR = 12.2, *P* = 0.018), confusion (OR = 91.95, *P* = 0.043), vertigo or dizziness (OR = 12.44, *P* = 0.042) and nausea (OR = 2183, *P* = 0.022).

**Figure 1.**
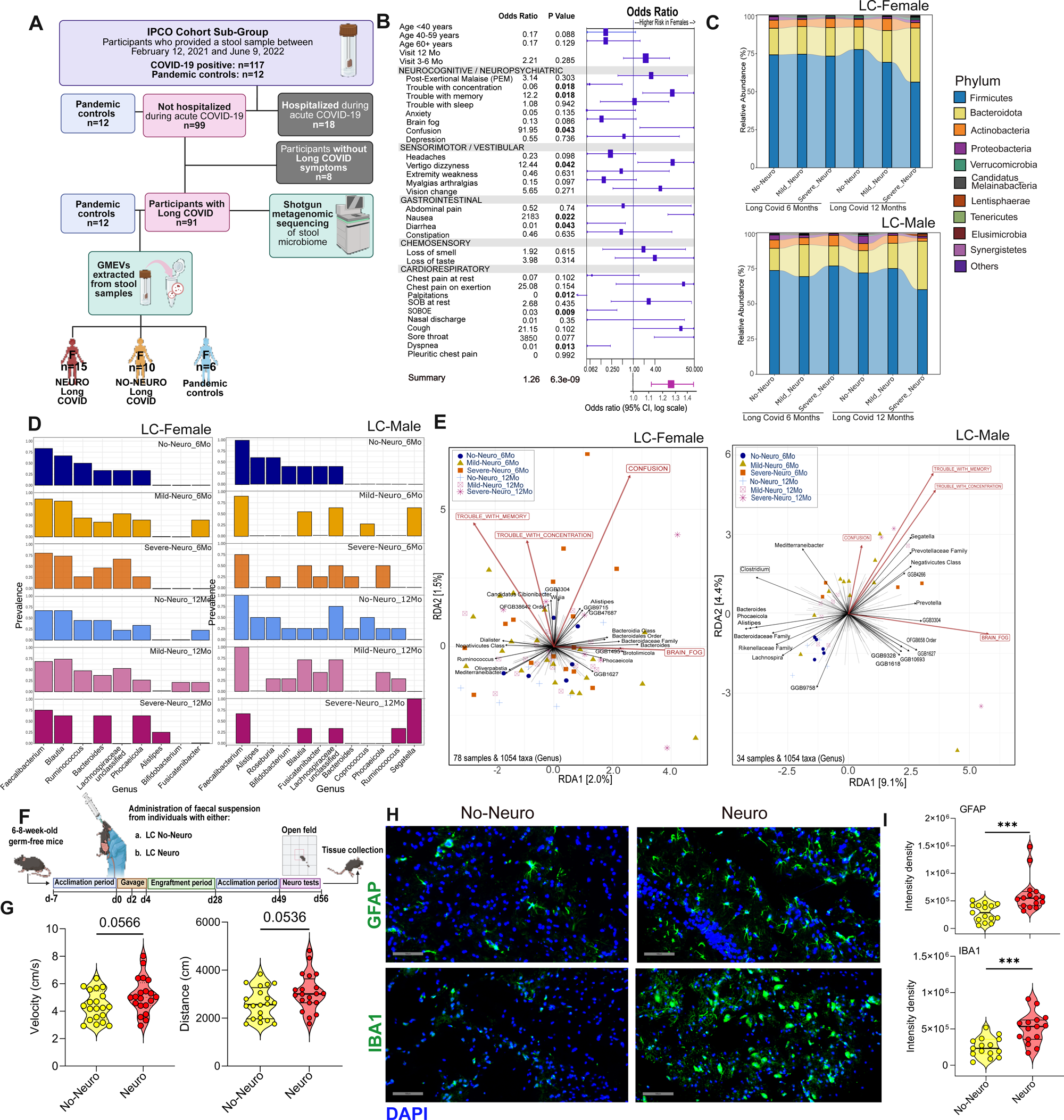
Sex-specific intestinal dysbiosis in long COVID associates with neurological symptom burden and transfers barrier and neuroinflammatory phenotypes to gnotobiotic mice. (A) IPCO cohort sub-study design and sample processing. Pandemic controls (PC; n = 12) and participants with long COVID (LC; n = 91; not hospitalized during acute infection) provided stool samples for shotgun metagenomic sequencing at 3-6 months and/or 12 months after infection; a subset was used for gut microbiota-derived extracellular vesicle (GMEV) extraction. Created in BioRender. Falcone, E. (2026) https://BioRender.com/g6cp3ke. (B) Forest plot showing odds ratios (ORs) for individual LC symptoms in females relative to males. Boxes indicate ORs and horizontal lines indicate 95% confidence intervals (log scale). (C) Phylum-level community composition in females (top) and males (bottom) stratified by neurological symptom burden and sampling time point (3-6 months and 12 months after infection). (D) Relative abundance of selected dominant bacterial genera in females (left) and males (right) stratified by neurological symptom burden and sampling time point. (E) Redundancy analysis (RDA) ordination of genus-level microbiome profiles in females (left) and males (right). Points represent samples colored by neurological symptom group and sampling time point; arrows (biplot vectors) indicate taxa and symptom loadings and their direction of association with the ordination axes. (F) Experimental design for fecal microbiota transfer into germ-free (GF) mice using stool from female LC donors with neurological symptoms (LC Neuro; n = 4 donors) or without neurological symptoms (LC No-Neuro; n = 4 donors). Created in BioRender. Aranguren, M. (2026) https://BioRender.com/8tx74nl. (G) Open-field test in GF mice colonized with LC No-Neuro or LC Neuro donor microbiota; mean velocity (left) and total distance travelled (right) are shown (each dot represents one mouse; n = 5 mice per group). Data are pooled from four independent donor experiments. (H) Representative immunofluorescence images of glial fibrillary acidic protein (GFAP; astrocytes; top) and ionized calcium-binding adaptor molecule 1 (IBA1; microglia; bottom) in brain sections from GF mice colonized with LC No-Neuro or LC Neuro donor microbiota (20×). Scale bar, 100 µm. (I) Quantification of GFAP and IBA1 fluorescence intensity density. Each dot represents one field (5 fields per mouse; n = 3 mice per group). Statistical analyses were performed using logistic regression for odds ratio estimation (B), Kruskal-Wallis test with Dunn’s post hoc correction for multi-group comparisons (C–E), permutational multivariate analysis of variance (PERMANOVA) for beta diversity analyses, and Wilcoxon rank-sum (two-sided) tests for two-group comparisons (G, I). Exact tests used are described in the Methods. Data are shown as medians with interquartile range unless otherwise indicated.

Across sexes, LC participants exhibited intestinal microbiome profiles that were distinct from PC participants at 3-6 months and 12 months following acute infection, with differences in alpha diversity, beta diversity and taxonomic features that persisted over time (Figure S1). To assess whether neurological symptom burden tracked with specific microbiome features, LC participants were stratified by neurological symptom burden. Female LC participants were categorized as having severe neurological symptoms (Severe Neuro; ≥3 of 8 assessed neurocognitive/neuropsychological symptoms), mild neurological symptoms (Mild Neuro; <3 symptoms), or no neurological symptoms (No-Neuro) (Figure 1B). In females, Severe Neuro LC participants showed higher microbial diversity and evenness than No Neuro LC participants (Figure S2A), together with shifts in community composition (Figure S2A). A similar pattern was observed in males, with neurological symptom burden associated with differences in diversity and beta diversity at 6 and/or 12 months (Figure S2C, D). Factor analysis of mixed data (FAMD) further supported an association of neurological and respiratory symptom dimensions with microbiome composition in both sexes (Figure S3). At the phylum level, community composition differed across neurological symptom strata and timepoints in both females and males (Figure 1C).

Taxonomic analyses identified sex-specific patterns of dysbiosis associated with neurological symptom burden. In females, Severe Neuro LC participants showed depletion in multiple genera within the class *Clostridia*, including *Ruminococcus*, *Lachnospiraceae,* and *Fusicatenibacter,* alongside enrichment of genus within order *Bacteroidales*, including *Bacteroides*, *Phocaeicola*, and *Alistipes*, relative to Mild Neuro and No-Neuro groups; these differences were maintained through 12 months (Figure 1D, left). In males, Severe and Mild Neuro LC participants showed depletion of *Alistipes*, *Roseburia* and *Bifidobacterium* relative to No-Neuro LC participants at both 6 and 12 months, with additional genus-level differences emerging at 12 months (Figure 1D, right). Ordination analyses linked these taxa shifts to specific neurological symptoms, including trouble with memory and concentration, confusion and brain fog, with stronger symptom-taxa associations in females (Figure 1E).

Given the higher burden of neurological symptoms among female participants, we tested whether the associated microbiota features were functionally transferable. Germ-free female mice were colonized with stool from female LC donors with neurological symptoms (LC Neuro; n = 3 donors) or from female LC donors without neurological symptoms (LC No Neuro; n = 3 donors) (Figure 1F). Mice colonized with LC Neuro microbiota showed evidence of impaired intestinal barrier integrity, including reduced/disrupted zonula occludens-1 (ZO-1) staining in colonic tissue (Figure S4B), and displayed neurobehavioral differences, including increased locomotion activity in the open-field test (Figure 1G, Figure S4A). These mice also developed neuroinflammatory changes, including accumulation of activated astrocytes in the hippocampal region and amoeboid-shaped activated microglia in the hindbrain (Figure 1H, I). In contrast, mice colonized with LC No-Neuro donor microbiota did not show these pathological features. Together, these data indicate that intestinal microbiota associated with neurological symptoms in female LC participants is sufficient to induce intestinal barrier dysfunction, neurobehavioral alteration and neuroinflammatory phenotypes in gnotobiotic mice.

### GMEVs from individuals with LC induce intestinal epithelial inflammation, impair barrier function and activate macrophages *in vitro*

To test whether GMEVs contribute to LC-associated mucosal and systemic inflammation, we isolated GMEVs from stool of LC participants with neurological symptoms (LC Neuro), LC participants without neurological symptoms (LC No-Neuro), and pandemic controls (PC). Transmission electron microscopy (TEM) and nanoparticle tracking analysis (NTA) confirmed vesicular morphology and broadly similar size distribution across groups, with comparable EV yield metrics as summarized in Figure S5.

We first assessed epithelial responses to GMEVs using induced pluripotent stem cell–derived human intestinal organoid (HIO) monolayers from healthy donors. Exposure to GMEVs (1μg/mL) increased inflammasome-associated transcripts in HIOs, including *NLRP3* and *IL1B*, relative to vehicle and PC-derived GMEVs, and increased *TNFSF13B* (encoding B-cell activating factor (BAFF)) (Figure 2A). At the protein level, LC-derived GMEVs promoted secretion of inflammatory mediators, with the most pronounced increases generally observed following exposure to LC Neuro GMEVs across measured cytokines and chemokines (including IL-1β, IL-8, TNF, CCL2 and CXCL10) (Figure 2A).

**Figure 2.**
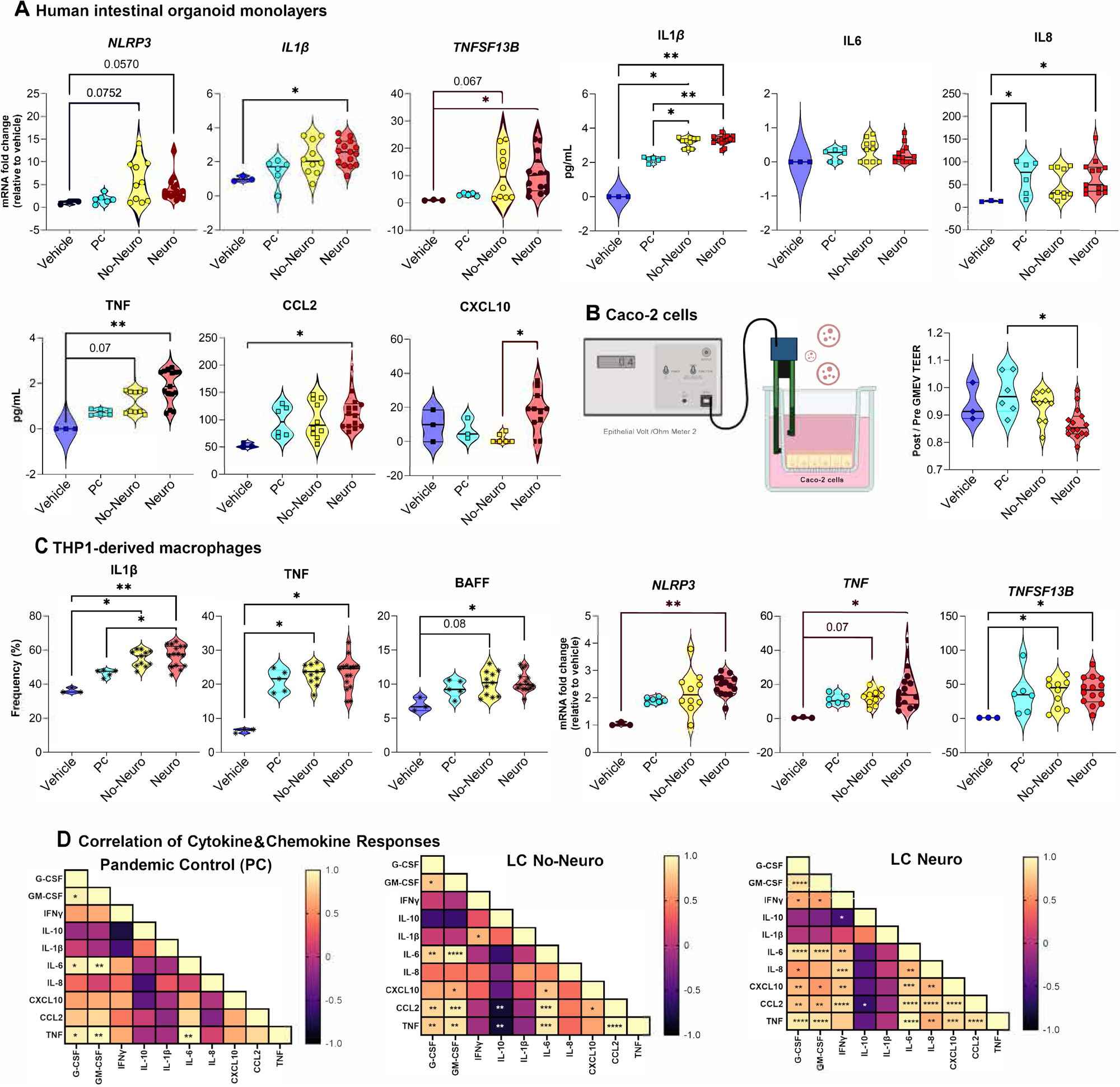
Gut microbiota-derived extracellular vesicles from individuals with long COVID induce epithelial inflammasome signaling, impair barrier function and activate macrophages *in vitro*. (A) Human induced pluripotent stem cell-derived intestinal organoid (HIO) monolayers were treated with vehicle (PBS; n = 3 independent experiments) or gut microbiota-derived extracellular vesicles (GMEVs; 1 μg/mL) isolated from pandemic controls (PC; n = 6 donors), long COVID (LC) donors without neurological symptoms (LC No-Neuro; n = 10 donors) or LC donors with neurological symptoms (LC Neuro; n = 15 donors). HIO transcript levels (including *NLRP3, IL1B, TNFSF13B*) were quantified by qRT-PCR at 5 h, and secreted mediators (including IL-1β, IL-6, IL-8, TNF, CCL2 and CXCL10) were quantified at 18 h using Meso Scale Discovery (MSD) assays. (B) Transepithelial electrical resistance (TEER) across Caco-2 monolayers measured 24 h after treatment with GMEVs (3μg/mL); values are shown as post/pre ratio. (C) THP-1-derived macrophages treated with vehicle (PBS; n = 3 independent experiments) or donor-derived GMEVs (PC, n = 6; LC No-Neuro, n = 10; LC Neuro, n = 15). Intracellular IL-1β and TNF frequencies and BAFF-associated responses were quantified by flow cytometry after 18 h stimulation with brefeldin A added for the final 5 h; macrophage inflammatory transcripts (including *NLRP3*, *TNF* and *TNFSF13B*) were measured by qRT–PCR at 5 h. (D) Pairwise correlation matrices of macrophage cytokine and chemokine responses across conditions (PC, LC No-Neuro and LC Neuro). Asterisks denote significant correlations (\**P* < 0.05, \*\**P* < 0.01, \*\*\**P* < 0.001, \*\*\*\**P* < 0.0001). In (A-C), each dot represents an independent donor-derived GMEV preparation (or an independent experiment for vehicle controls); center lines indicate medians and boxes indicate interquartile ranges. Statistical analyses were performed using Kruskal-Wallis test with Dunn’s post hoc correction for multi-group comparisons. Correlations were assessed using Spearman’s two-sided test.

Increased intestinal permeability and microbial translocation are hallmarks of chronic intestinal inflammation^22,24^, and individuals with LC exhibit elevated circulating markers of barrier dysfunction and microbial translocation, including soluble zonulin, lipopolysaccharide-binding protein (LBP) and β-D-glucan^32,41^. We therefore evaluated whether GMEVs directly alter epithelial barrier integrity by measuring trans-epithelial electrical resistance (TEER) across Caco-2 cell monolayers. LC Neuro GMEVs produced the strongest reduction in TEER (post-pre), consistent with impaired barrier function, whereas LC No-Neuro GMEVs exerted more modest effects and PC-derived GMEVs had minimal impact (Figure 2B).

We next asked whether GMEVs promote innate immune activation that could plausibly contribute to systemic inflammation. THP-1-derived macrophages responded to GMEVs from all donor groups compared to vehicle, consistent with a conserved immunostimulatory capacity of microbiota-derived vesicles (Figure 2C). However, LC Neuro GMEVs elicited the most robust macrophage activation, with higher frequencies and/or intensities of IL-1β-, TNF- and BAFF-associated responses compared with LC No-Neuro and PC conditions (Figure 2C; Figure S6D). In parallel, LC-derived GMEVs increased expression of inflammatory transcripts in macrophages, including *NLRP3*, *TNF* and *TNFSF13B*, with the strongest responses again observed in the LC Neuro condition (Figure 2C).

Finally, to compare inflammatory network structure across donor groups, we examined cytokine/chemokine relationships induced by GMEVs. Correlation analyses revealed a denser pattern of coordinated mediator responses in macrophages stimulated with LC Neuro GMEVs than with LC No-Neuro or PC GMEVs, including stronger coupling between *TNFSF13B* and multiple inflammatory readouts (Figure S6A-C).

Together, these data indicate that LC-derived GMEVs are sufficient to trigger epithelial inflammatory programs and macrophage activation *in vitro*, and that GMEVs from LC Neuro donors most consistently associate with barrier-disruptive activity and coordinated multi-cytokine inflammatory responses.

### Extracellular vesicles derived from the gut microbiota of individuals with LC induce a pro-inflammatory response in iPSC-derived microglia

Because transplantation of LC-associated microbiota induced neuroinflammatory phenotypes in germ-free mice and LC-derived GMEVs activated macrophages *in vitro*, we asked whether GMEVs can also directly modulate central nervous system innate immune cells. We therefore exposed induced pluripotent stem cell-derived microglia (iMGLs) to donor-derived GMEVs from LC participants (LC Neuro and LC No-Neuro) or pandemic controls (PC) and profiled transcriptional responses by RNA sequencing. Global ordination separated iMGLs stimulated with LC-derived GMEVs from those treated with PC-derived GMEVs, with LC Neuro and LC No-Neuro conditions clustering together, indicating broadly similar microglial responses across LC donor subgroups (Figure 3A). Differential expression analysis identified 2,631 genes altered in iMGLs exposed to LC-derived versus PC-derived GMEVs (adjusted *P* < 0.05) (Figure 3B). Gene ontology enrichment highlighted a coordinated activation of immune and inflammatory processes, including innate immune response programs, chemotaxis-related terms and cytokine-linked pathways (Figure 3C; Figure S7A). Consistent with this, pathway-level analyses identified significant enrichment of immune and infection -associated signatures, ranked among the top differentially regulated pathways, including the Kyoto Encyclopedia of Genes and Genomes (KEGG) “COVID-19” pathway (Figure 3D; Figure S7B, C). Gene set enrichment analysis further showed preferential induction of inflammatory response modules in LC-GMEV-treated iMGLs, including interferon- and cytokine-associated signaling and neutrophil-linked programs (Figure 3E). In addition, enrichment analyses identified pathways with established roles in neuroimmune communication, including axon guidance, which showed broad upregulation of constituent genes in iMGLs treated with LC-derived GMEVs (Figure 3F). Protein-protein interaction network analysis of highly induced genes revealed interconnected modules centered on inflammatory and myeloid activation nodes, including networks annotated for regulation of neuroinflammatory responses (Figure S8A).

**Figure 3.**
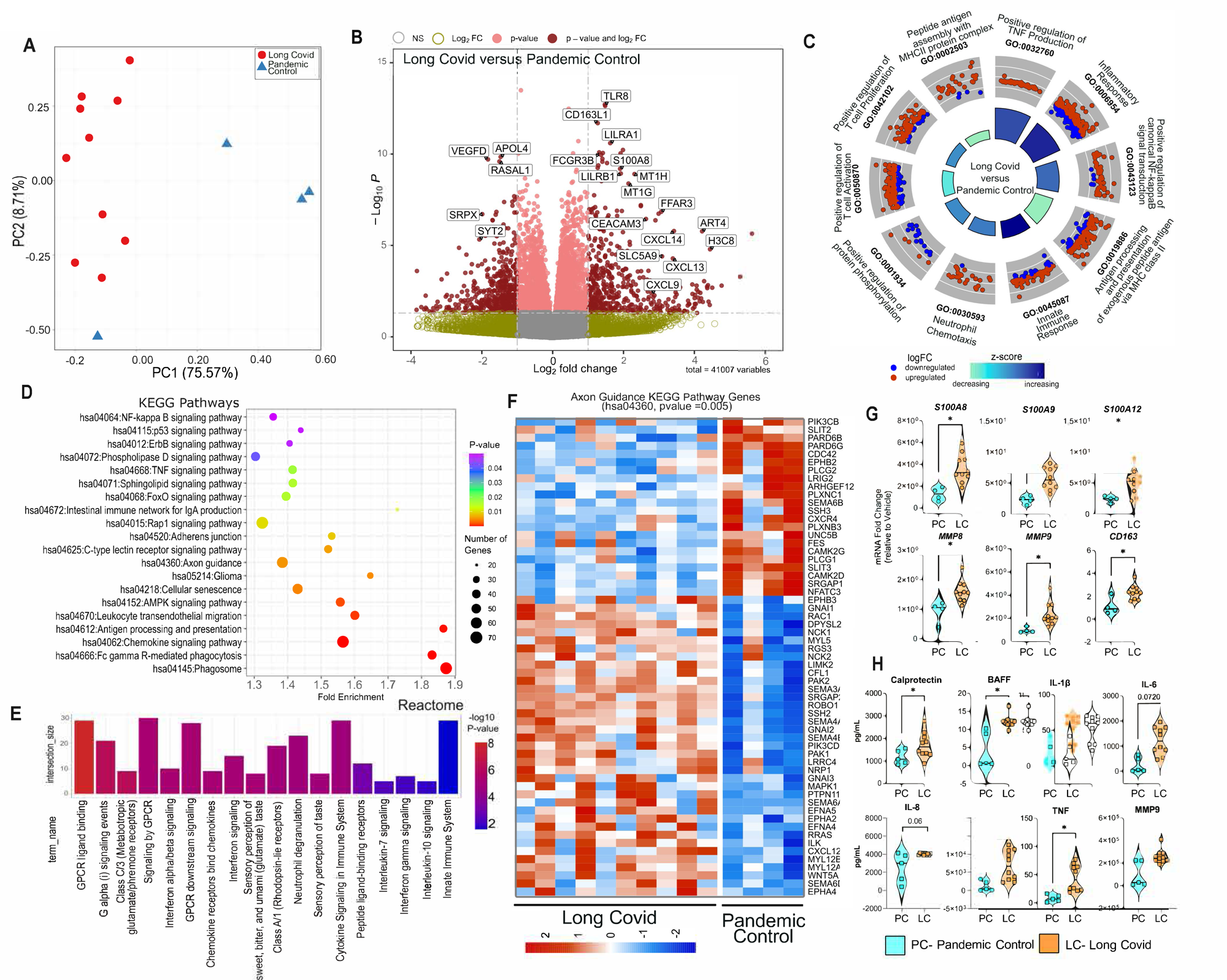
Gut microbiota-derived extracellular vesicles from individuals with long COVID induce a neuroinflammatory transcriptional program in induced pluripotent stem cell-derived microglia. Induced pluripotent stem cell-derived microglia (iMGLs) were exposed for 16 h to gut microbiota-derived extracellular vesicles (GMEVs; 1 µg/mL) from long COVID donors (LC; n = 10), pandemic controls (PC; n = 4), or vehicle (PBS; n = 3 independent experiments), followed by bulk RNA sequencing. (A) Principal component analysis (PCA) of transcriptomes from GMEV-treated iMGLs. Percent variance explained by each principal component is indicated on the axes. (B) Volcano plot of differentially expressed genes in LC versus PC conditions (adjusted *P* < 0.05; DESeq2, Benjamini-Hochberg correction)). Selected significantly regulated genes are annotated. (C) Gene Ontology (GO) circle plot summarizing enriched biological process terms among differentially expressed genes. Bar height indicates significance (−log10 adjusted *P* value), color indicates Z-score (directionality), and scatter points represent individual genes plotted by log2 fold change. (D) Top 20 enriched KEGG pathways ranked by −log10 adjusted *P* value. (E) Reactome pathway enrichment analysis highlighting immune and cytokine signaling pathways among LC-induced genes. (F) Heatmap of genes within the KEGG axon guidance pathway (hsa04360; adjusted *P* = 0.005). Rows represent genes and columns represent samples; red indicates upregulation and blue indicates downregulation (Z-score scaled). (G) Quantitative RT-PCR validation of selected inflammatory and neuroimmune-associated transcripts in iMGLs treated with vehicle, PC-derived GMEVs or LC-derived GMEVs. (H) Cytokine and chemokine secretion measured by Meso Scale Discovery (MSD) assays following GMEV stimulation. In (G, H) each dot represents one donor-derived GMEV preparation (LC or PC) or one independent vehicle experiment; center lines indicate medians and boxes indicate interquartile ranges. Statistical analyses for RNA sequencing were performed using DESeq2 with Benjamini-Hochberg correction for multiple testing. For (G, H), multi-group comparisons were performed using Kruskal-Wallis test with Dunn’s post hoc correction.

We validated key transcriptomic changes by targeted qPCR, confirming induction of inflammatory and myeloid-associated genes (for example, *IL1B*, *TNF*, *IL6*, *AIF1* and neutrophil-linked S100 family transcripts) in iMGLs exposed to LC-derived GMEVs compared with PC-derived GMEVs and vehicle (Figure 3G; Figure S8B). At the protein level, LC-derived GMEVs increased secretion of inflammatory mediators, including IL-6, TNF and CXCL1 (Figure 3H). Notably, *TNFSF13B* induction and increased BAFF protein were also observed, paralleling epithelial and macrophage responses and identifying BAFF as a convergent inflammatory mediator in response to GMEVs across multiple cellular compartments (Figure 3G, H).

Together, these data show that GMEVs from individuals with LC, irrespective of neurological symptom subgroup, are sufficient to elicit a robust human microglial activation state marked by coordinated induction of inflammatory and neuroimmune-associated gene programs, thereby supporting a role for LC-associated GMEVs in shaping microglial responses that may contribute to neuroinflammation in LC.

### Oral administration of GMEVs from individuals with LC remodels the intestinal microbiota in wild-type mice

We next asked whether GMEVs are sufficient to reshape intestinal microbial communities *in vivo*. Wild-type (WT) mice received oral gavage of donor-derived GMEVs (5μg/mL) from LC Neuro participants, LC No-Neuro participants or PCs, or vehicle alone, 3 times per week for 6 weeks. To assess whether orally administered vesicles disseminate beyond the gut, we tracked fluorescently labelled bacterial EVs (derived from *Escherichia coli* as a surrogate for GMEVs). Eight hours after gavage, fluorescent signal was detected in the gastrointestinal tract and in peripheral organs, including liver and kidney, and was also measurable in brain tissue, whereas signal was not detected in mice receiving unlabeled vesicles (Figure S9).

To determine whether GMEVs alter intestinal community structure, we performed 16S rRNA gene sequencing on fecal samples collected after 6 weeks of treatment. Relative to vehicle and PC groups, administration of GMEVs from LC donors, irrespective of neurological symptom status, was associated with reduced alpha diversity, with decreases in Shannon diversity and Pielou’s evenness indices (Figure 4A). At the phylum level, mice receiving GMEVs from LC Neuro donors showed the most pronounced compositional shifts, including altered relative abundance of Bacillota to Bacteroidota and reductions in Actinomycetota and Verrucomicrobiota compared to controls groups (Figure 4B). Principal coordinate analysis based on Bray-Curtis dissimilarity showed clear separation of microbiota profiles in mice treated with LC-derived GMEVs from those receiving vehicle or PC-derived GMEVs (Figure 4C). Notably, mice administered GMEVs from LC No-Neuro donors displayed a bifurcated clustering pattern, with partial overlap with both control and LC Neuro clusters, consistent with heterogeneous responses to LC No-Neuro GMEVs.

**Figure 4.**
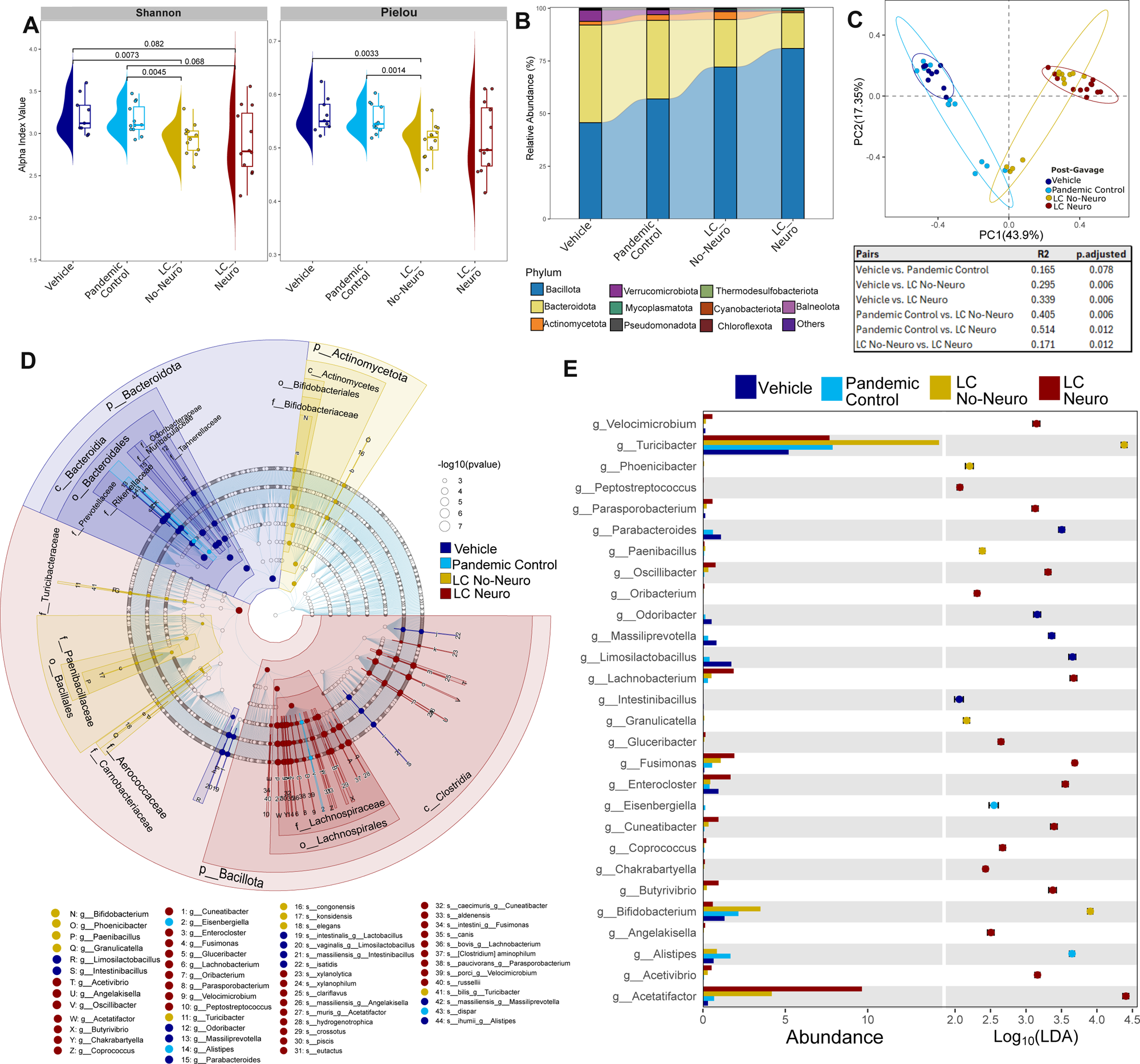
Oral administration of gut microbiota-derived extracellular vesicles from individuals with long COVID remodels the fecal microbiota of wild-type mice. Female C57BL/6J mice (6-8 weeks old) received oral gavage of vehicle (PBS; n = 9 mice total; 3 mice per run across 3 independent experimental runs) or gut microbiota-derived extracellular vesicles (GMEVs; 5 µg per dose) isolated from pandemic controls (PC; 3 GMEV donors; n = 4 mice per donor), LC donors without neurological symptoms (LC No-Neuro; 3 GMEV donors; n = 4 mice per donor) or LC donors with neurological symptoms (LC Neuro; 3 GMEV donors; n = 4 mice per donor). GMEVs were administered 3 times per week for 6 weeks, after which fecal microbiota composition was profiled by 16S rRNA gene sequencing. The experiment was performed in 3 independent runs, each using an independent donor-derived GMEV preparation per group. (A) Alpha diversity (Shannon diversity and Pielou’s evenness). Multi-group comparisons were performed using Kruskal-Wallis test with Dunn’s post hoc correction. (B) Relative abundance of bacterial phyla across groups. (C) Principal coordinates analysis (PCoA) of Bray-Curtis dissimilarity showing separation of fecal communities by treatment group. Group differences were assessed using PERMANOVA. (D) Cladogram from linear discriminant analysis effect size (LEfSe) showing taxa that differ in relative abundance across groups (Kruskal-Wallis test, α = 0.05; linear discriminant analysis (LDA) score ≥ 2). (E) LEfSe LDA scores for differentially abundant genera; bars indicate effect size and points indicate relative abundance.

We next identified taxa contributing to these treatment-associated differences. Linear discriminant analysis effect size (LEfSe) revealed distinct microbial features associated with each treatment group (Figure 4D). Mice receiving LC Neuro-derived GMEVs were enriched for taxa within the family Lachinospiraceae and the class *Clostridia*, including the genera *Acetatifactor, Lachnobacterium, Fusimonas, Enterocloster, and Velocimicrobium* (Figure 4E). In contrast, treatment with LC No-Neuro-derived GMEVs was associated with enrichment of a partially distinct set of genera, including *Turicibacter* and *Bifidobacterium* (Figure 4E).

Together, these data indicate that oral exposure to LC-derived GMEVs is sufficient to remodel the fecal microbiota in WT mice, with LC Neuro-derived GMEVs associated with the most distinct compositional shifts.

### Oral administration of GMEVs from individuals with LC induces intestinal inflammation, neurobehavioral alterations and neuroinflammation *in vivo*

We next tested whether oral exposure to donor-derived GMEVs is sufficient to elicit intestinal and systemic inflammatory changes accompanied by neurobehavioral and neuroinflammatory phenotypes in wild-type mice. Female C57BL/6J mice received oral gavage of GMEVs (5 µg per dose) isolated from LC donors with neurological symptoms (Neuro), LC donors without neurological symptoms (No-Neuro) or pandemic controls (PC), or vehicle alone, 3 times per week for 6 weeks, followed by neurobehavioral testing and tissue collection (Figure 5A).

**Figure 5.**
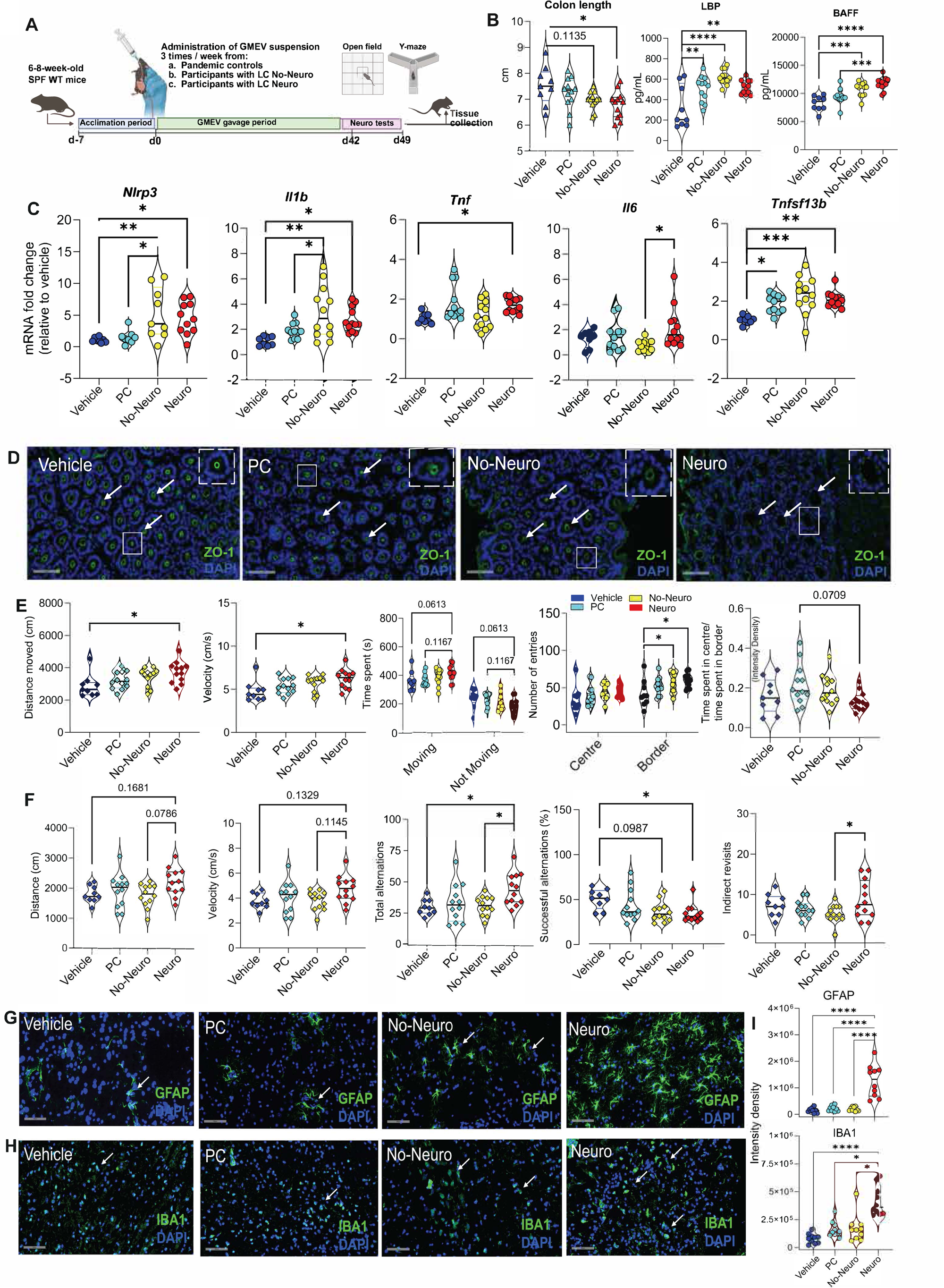
Oral administration of gut microbiota-derived extracellular vesicles from individuals with long COVID induces intestinal inflammatory signatures, behavioral alterations and glial activation in wild-type mice. Female C57BL/6J mice (6-8 weeks old) received oral gavage of gut microbiota–derived extracellular vesicles (GMEVs; 5 µg per dose) isolated from LC donors without neurological symptoms (No-Neuro; 3 GMEV donors; n = 4 mice per donor), LC donors with neurological symptoms (Neuro; 3 GMEV donors; n = 4 mice per donor) or pandemic controls (PC; 3 GMEV donors; n = 4 mice per GMEV donor), or vehicle (PBS; n = 9 mice total; 3 mice per run across 3 independent experimental runs), 3 times per week for 6 weeks. The experiment was performed in 3 independent runs using one donor-derived GMEV preparation per group per run. (A) Experimental design. Created in BioRender. Falcone, E. (2026) https://BioRender.com/nk3kim1. (B) Colon length and serum levels of lipopolysaccharide-binding protein (LBP) and B-cell activating factor (BAFF) at endpoint. (C) Colonic epithelial expression of *Nlrp3*, *Il1b*, *Tnfsf13b*, *Tnf* and *Il6* measured by qRT-PCR. (D) Representative immunofluorescence images of zonula occludens-1 (ZO-1; green) in colonic sections (green) with DAPI nuclear counterstain (blue); dashed boxes indicate regions shown at higher magnification (40×). Scale bar, 100 µm. Representative images are shown from n = 2 mice per group. (E) Open-field test, including distance travelled, velocity, time spent moving versus not moving and center-border entries, and time spent in center relative to border. (F) Y-maze spontaneous alternation behavior, including distance travelled, velocity, total alternations, proportion of successful alternations, and indirect revisits. (G, H) Representative immunofluorescence images of glial fibrillary acidic protein (GFAP; astrocytes; G) and ionized calcium-binding adapter molecule 1 (IBA1; microglia; H) in brain sections, with DAPI counterstain. **(I**) Quantification of GFAP and IBA1 fluorescence intensity density. Each dot represents one field (5 fields per mouse; n=3 mice per group). In (B, C, E, F, I), multi-group comparisons were performed using Kruskal-Wallis test with Dunn’s post hoc correction. Each dot represents one mouse unless otherwise indicated. Violin plots show distribution; center lines indicate medians and boxes indicate interquartile ranges.

Mice receiving LC-derived GMEVs showed evidence of intestinal and systemic inflammatory activation. Colon length was modestly reduced in GMEV-treated groups compared with vehicle, with the largest reduction observed in the Neuro GMEV group (Figure 5B). Circulating markers were also altered as serum lipopolysaccharide-binding protein (LBP) was increased in GMEV-treated groups relative to vehicle, and serum BAFF was significantly elevated in mice receiving Neuro-derived GMEVs compared with controls (Figure 5B). In colonic epithelium, Neuro-derived GMEVs increased expression of inflammasome- and inflammatory-associated genes, including *Nlrp3*, *Il1b* and *Tnfsf13b*, relative to controls, whereas changes in *Tnf* and *Il6* were more modest (Figure 5C). Consistent with barrier perturbation, ZO-1 immunostaining showed more frequent junctional discontinuities in mice receiving LC-derived GMEVs, which was most apparent in the Neuro GMEV group, relative to vehicle and PC conditions (Figure 5D).

We next evaluated whether these inflammatory changes were accompanied by alterations in behavior. Anxiety-like behavior was evaluated using the open-field test, and short-term spatial memory was assessed using the Y-maze^42^. In the open-field test, mice receiving Neuro-derived GMEVs travelled a greater distance and displayed higher mean velocity than control groups, indicating increased locomotor activity (Figure 5E). Center-border measures showed more modest differences as time spent in the border zone increased in the Neuro group, whereas center time and the center-to-border time ratio showed non-significant trends (Figure 5E), a pattern that can be suggestive of anxiety-like behavior but is not definitive in isolation. In the Y-maze, Neuro-derived GMEVs increased total alternations but reduced the proportion of successful alternations, with a concurrent increase in indirect revisits, consistent with altered spontaneous alternation behavior (Figure 5F) and suggestive of impaired short-term spatial working memory. Memory impairment is a common symptom reported by LC Neuro participants in the IPCO cohort (Figure 1B).

Finally, building on the behavioral phenotypes and the observation that orally administered bacterial EVs can be detected beyond the gut (Figure S9), we assessed glial activation in brain tissue collected after behavioral testing. Mice treated with Neuro-derived GMEVs showed increased GFAP immunoreactivity, with prominent signal in regions proximal to the hippocampus, and increased IBA1-positive microglial signal with an activated morphology in the hindbrain, compared with control groups (Figure 5G-I). These findings are consistent with induction of neuroinflammatory changes following oral exposure to LC-derived vesicles.

Together, these data show that oral administration of GMEVs derived from individuals with LC, most consistently for Neuro-derived preparations, is sufficient to induce intestinal inflammatory responses and barrier perturbation, accompanied by systemic inflammatory signatures, neurobehavioral alterations, and glial activation *in vivo*. These findings implicate GMEVs as functional mediators linking LC-associated dysbiosis to gut-brain axis perturbations that may contribute to neurological sequelae in LC.

## Discussion

Persistent symptoms following viral infection are increasingly linked to alterations in the intestinal microbiota^25,26,43^, yet the molecular intermediates that enable dysbiotic communities to influence distal host tissues remain poorly defined. Here, we identify GMEVs as functional mediators of host-microbe communication in LC. By integrating longitudinal human microbiota profiling with gnotobiotic transfer, mechanistic epithelial and myeloid models, and *in vivo* vesicle administration, our findings establish GMEVs as biologically active mediators of inter-kingdom communication capable of reshaping mucosal, systemic, and neuroimmune responses.

Consistent with previous reports in acute and post-acute syndromes^17–20^, we observed sustained intestinal dysbiosis in individuals with LC that persisted for at least 12 months after infection. Importantly, microbiome alterations correlated with prototypical LC symptoms, particularly neurological manifestations, reinforcing a link between gut microbial composition and disease phenotype. Transfer of LC-associated microbiota into germ-free mice recapitulated features of intestinal barrier dysfunction, systemic inflammation, and neuroinflammation, with effects most pronounced in mice colonized with microbiota from individuals with neurological symptoms. These findings extend prior fecal microbiota transplantation studies^44^ by demonstrating symptom-severity-dependent effects and support a contributory role for the gut microbiota in LC pathophysiology.

Microbial extracellular vesicles represent a fundamental mode of host-microbe communication, yet their role in shaping host inflammatory tone following viral perturbation of the microbiota has remained poorly defined. Although EVs derived from host cells have been implicated in acute COVID-19 and LC^45,46^, our data show that microbiota-derived EVs alone are sufficient to induce LC-relevant phenotypes. GMEVs from LC donors promoted inflammasome activation, epithelial barrier disruption, and inflammatory cytokine and chemokine production in intestinal epithelial cells and macrophages, and induced a coordinated inflammatory transcriptional program in human iPSC-derived microglia. Notably, while GMEVs from LC Neuro and LC No-Neuro donors elicited broadly similar microglial transcriptomic responses, Neuro-derived GMEVs consistently induced stronger epithelial and macrophage activation, suggesting cell-type-specific sensitivity to GMEV-associated inflammatory signals.

*In vivo*, chronic oral administration of LC-derived GMEVs was sufficient to alter gut microbial composition, disrupt intestinal barrier integrity and induce systemic inflammation, recapitulating several features observed in our microbiota transfer model. Fluorescent tracing experiments demonstrated systemic dissemination of orally administered bacterial EVs, with detectable signal in peripheral organs and brain tissue, consistent with previous reports^47^. Mice receiving GMEVs from LC Neuro donors developed increased locomotor activity, altered exploratory behavior and impaired spontaneous alternation performance, accompanied by accumulation of GFAP-positive reactive astrocytes and IBA1 positive activated microglia in discrete brain regions. These findings indicate that LC-derived GMEVs induce spatially localized glial activation consistent with neuroinflammatory processes that may not be captured by bulk tissue analyses.

Among induced inflammatory pathways, BAFF-associated signaling emerged as a component of a broader vesicle-driven response network. Elevated BAFF levels tracked with LC severity in our cohort and were reproducibly induced by LC-derived GMEVs across epithelial, myeloid, and glial systems *in vitro* and *in vivo*. Excess BAFF is a recognized driver of chronic immune activation, B-cell dysregulation, and autoantibody production^48–53^, and increased BAFF signaling has been reported in chronic inflammatory states characterized by sustained immune activation and intestinal barrier dysfunction, including HIV infection^48,49,54–56^. In our system, BAFF induction likely reflects amplification of vesicle-triggered innate immune activation, with downstream consequences for B-cell homeostasis and adaptive immune regulation, positioning BAFF as a downstream effector within a GMEV-driven inflammatory cascade linking dysbiosis to systemic immune dysregulation.

Mechanistically, GMEVs likely engage host cells through multiple pathways, including pattern-recognition receptor signaling triggered by surface-associated microbial components, vesicle internalization, and delivery of biologically active cargo. In this framework, vesicles function as concentrated, protected carriers of microbial cargo, enabling coordinated delivery of immune-modulatory signals beyond the intestinal lumen. Differences in EV composition between donor groups, potentially reflecting microbial taxonomic shifts or structural variation in microbial products such as lipopolysaccharide^57,58^, may underlie the graded inflammatory responses observed across GMEVs derived from different LC donor groups. Although LC is undoubtedly multifactorial, our data support a model in which GMEVs amplify and perpetuate inflammatory signaling downstream of intestinal dysbiosis.

Several limitations should be considered. Although the IPCO cohort is longitudinal and clinically characterized, LC is heterogeneous and residual confounding by factors that influence the microbiome, including diet, medication exposures and co-morbidities, cannot be fully excluded. The biodistribution experiment used fluorescently labelled *Escherichia coli* EVs as a surrogate for complex GMEVs. Thus, the kinetics and tissue tropism of human donor-derived vesicles may differ. Neurobehavioral assays provide operational measures of exploration and spontaneous alternation but do not uniquely map to specific neuropsychiatric constructs. Accordingly, the observed open-field and Y-maze changes should be interpreted as behavioral alterations consistent with neuroimmune perturbation rather than definitive measures of anxiety or memory impairment. Finally, the precise microbial sources and vesicle cargo responsible for host activation were not resolved. Defining the molecular determinants of GMEV bioactivity and the host pathways required for their effects will be important for therapeutic translation.

Together, these findings identify GMEVs as functional effectors linking dysbiotic microbial communities to mucosal, systemic, and neuroimmune inflammation. By integrating human cohort analyses with mechanistic *in vitro* and *in vivo* models, this work advances a vesicle-centered framework for host-microbe communication in post-infectious inflammatory states and potentially other conditions characterized by microbiota-driven immune dysregulation. Targeting vesicle production, cargo composition, or host sensing pathways may therefore represent therapeutic strategies to modulate persistent inflammation following infection.

## Methods

### Study design and population

In response to the COVID-19 pandemic, we established the *Institut de Recherches Cliniques de Montréal* (IRCM) Post-COVID-19 (IPCO) research clinic, which integrates clinical care into a prospective observational cohort study with an associated biobank (IPCO protocol #2021-1092, ClinicalTrials.gov: <u>NCT04736732</u>). Participants are followed longitudinally for up to 24 months with standardized clinical assessments and biospecimen collection. Adults (≥18 years) with confirmed SARS-CoV-2 infection at least 3 months before enrolment and persistent symptoms not attributable to alternative diagnoses were recruited as Long COVID (LC) participants. Pandemic controls (PC) were individuals without persistent symptoms and without reported SARS-CoV-2 infection or positive testing. Participants underwent in-person visits at enrolment and at 6, 12 and 24 months after infection, with a telephone follow-up at 18 months. Baseline demographic and clinical characteristics of participants included in microbiome analyses are provided in Table S1.

### Symptom definitions and neurological stratification

Participants reported symptoms using standardized questionnaires administered at each visit. For analyses stratifying LC participants by neurological symptom burden, 8 self-reported symptoms were considered: post-exertional malaise, trouble with concentration, trouble with memory, trouble with sleep, anxiety, brain fog, confusion and depression. For binary classification used in downstream functional experiments (microbiota transfer and GMEV studies), participants reporting ≥1 of these symptoms were classified as LC Neuro, whereas those reporting none were classified as LC No-Neuro. For microbiome association analyses assessing symptom burden (Figure 1), LC participants reporting neurological symptoms were further stratified into Mild Neuro (<3 symptoms) or Severe Neuro (≥3 symptoms) groups to evaluate dose-response relationships between neurological symptom burden and microbiome composition.

### Sample collection and processing

At each study visit, participants provided stool, blood, saliva, and urine samples. Serum and plasma were isolated by centrifugation. Peripheral blood mononuclear cells (PBMCs) were isolated using SepMate tubes (STEMCELL Technologies). All biospecimens were aliquoted and stored at −80°C, and PBMCs were cryopreserved in liquid nitrogen.

### Stool DNA extraction and microbiome sequencing

#### Human samples

Stool DNA was extracted using the QIAamp PowerFecal Pro DNA Kit (QIAGEN) with mechanical homogenization. Shotgun metagenomic libraries were prepared after quality control and sequenced on an Illumina NovaSeq 6000 platform (CosmosID). Host reads were removed by alignment to the human reference genome (GRCh38). Read quality was assessed using FastQC^59^. Functional profiling was performed using HUMAnM (v3.9)^60^ and taxonomic profiling using MetaPhlAn (v4.1.1)^61^.

Microbiome analyses were performed in R (v4.2) using phyloseq (v1.48.0)^62^ and tidyverse packages^63^. Alpha diversity was assessed using the Simpson index Pielou’s evenness, and beta diversity using Bray-Curtis dissimilarity with principal coordinate analysis (PCoA). Differentially abundant taxa were identified using Linear Discriminant Analysis (LDA) Effect Size (LEfSe)-based approaches implemented in edgeR^64^ and microbiomeMarker^65^, with a Kruskal-Wallis significance threshold of *P* ≤ 0.05 and an LDA score cutoff as indicated in the corresponding figure legends.

#### Mouse stool 16S rRNA sequencing

Mouse fecal pellets were collected into sterile tubes, snap-frozen on dry ice and stored at −80°C. DNA was extracted using the DNeasy PowerSoil Pro QIAcube HT Kit (QIAGEN). The V4 region of the 16S rRNA gene was amplified and sequenced on an Illumina MiSeq platform (McGill Center for Microbiome Research). Read quality was assessed using FastQC^59^, adapters and primers were trimmed with Trimmomatic^66^. Paired-end reads were merged with PANDAseq^67^ and chimeras removed as described previously^68^. Sequences were clustered using CD-HIT-EST^69^ and taxonomically classified using Kraken2^70^ against an in-house RefSeq database (updated November 2023).

Alpha diversity was assessed using Shannon diversity and Pielou’s evenness. Beta diversity was evaluated using Bray-Curtis dissimilarity and visualized by PCoA. Group differences were assessed by permutational multivariate analysis of variance (PERMANOVA; adonis2, vegan v2.6-6.1; 1000 permutations)^71^, with pairwise comparisons adjusted for multiple testing as indicated. Differentially abundant taxa were identified using LEfSe (edgeR^64^ and microbiomeMarker^65^; LDA cutoff and significance thresholds as indicated in the corresponding figure legends).

### Animals and housing

Wild-type (WT) C57BL/6J female mice (6-8 weeks old) were obtained from The Jackson Laboratory and housed under specific pathogen-free conditions at the IRCM animal facility with a 12-hour light/dark cycle with controlled temperature and humidity and ad libitum access to food and water. Germ-free (GF) C57BL/6 female mice (5-7 weeks old) were obtained from the Germ-Free and Gnotobiotic Platform at the University of Calgary (K. McCoy) and maintained in flexible film isolators (Class Biologically Clean) under sterile conditions at the IRCM gnotobiotic facility. Mice were routinely screened for contamination. All animal procedures were approved by the IRCM Animal Care Committee.

### Fecal microbiota transplantation into germ-free mice

Mouse experimental group sizes and anonymized donor sample identifiers are summarized in Table S2. GF mice were orally gavaged 3 times per week with 200 μl of a stool suspension prepared by resuspending 200mg of human donor stool in 1 mL sterile PBS. After the first gavage, mice were transferred to positive-pressure cages to prevent cross-contamination. Microbiota were allowed to engraft for 3 weeks before mice were transferred to irradiated racks for acclimation, behavioral testing and tissue collection.

#### GMEV isolation and characterization

Gut microbiota-derived extracellular vesicles (GMEVs) were isolated from human stool by size-exclusion chromatography. Stool suspensions were sequentially centrifuged and filtered (0.45 μm and 0.22 μm) to remove bacteria and debris, concentrated by ultrafiltration (100 kDa cut-off), and fractionated using qEV Original 35 nm Gen 2 column (IZON). GMEV-enriched fractions were pooled, aliquoted, and stored at −80°C. Particle size and concentration were assessed by nanoparticle tracking analysis (ZetaView PMX120). Vesicular morphology was confirmed by transmission electron microscopy (Tecnai G2 Spirit Twin). Protein concentration was measured using the Pierce Bovine Serum Albumin Protein assay (Thermo Fisher Scientific).

### Neurobehavioral testing

Neurobehavioral testing was performed after microbiota engraftment (GF colonization experiments) or after the oral gavage regimen (WT GMEV experiments), as indicated in the relevant figure legends. Testing was recorded using the EthoVision and analyzed blinded to group allocation where feasible.

#### Y-maze spontaneous alternation

The Y-maze test was used to assess spontaneous alternation behavior, which is sensitive to short-term spatial working memory^42^. Mice were placed in the center of a 3-armed maze and allowed to explore freely for 8 minutes. Arm entries were scored by EthoVision. Consecutive triplets of arm entries were defined as alternations; triplets comprising 3 different arms were scored as successful alternations. Performance was calculated as the ratio of successful alternations to total alternations, with direct and indirect revisits also quantified. A lux meter was used to ensure that illumination was standardized across all 3 arms (approximately 150 lux).

#### Open-field test

Exploration and anxiety-like behavior were assessed using the open-field test^72^. Mice were placed in the arena and recorded by EthoVision. Time spent in the center versus periphery (border), the number of center-border transitions, total distance travelled, and velocity were quantified. A lux meter was used to ensure that illumination was standardized across the whole arena (approximately 150 lux).

### Oral gavage of GMEVs in wild-type mice

Group sizes and anonymized donor sample identifiers are provided in Table S2. Female WT C57BL/6J mice (6-8 weeks old; The Jackson Laboratory), maintained under specific pathogen-free conditions, received GMEVs by oral gavage (5 μg per dose in 200 μl sterile PBS) or vehicle (sterile PBS) 3 times per week for 6 weeks. Following the treatment period, mice underwent behavioral testing prior to euthanasia and tissue collection. Colon length was measured at necropsy. Colon and brain tissue were processed for histology and molecular analyses as described below.

### Biodistribution of orally administered EVs

To assess biodistribution following oral delivery, extracellular vesicles were isolated from Escherichia coli (ATCC 25922) culture supernatants by ultrafiltration and size-exclusion chromatography (qEV 35 nm Gen 2; Izon). Vesicles were labelled with Vybrant DiD (Invitrogen) according to the manufacturer’s instructions and administered to female BALB/c mice by oral gavage (10 µg dose of EVs). Fluorescence was acquired 8h after gavage using a Xenogen IVIS 200 system (PerkinElmer) under isoflurane anesthesia. Mice were then euthanized, and organs were harvested for ex vivo fluorescence imaging.

### Blood collection and serum isolation

Mice were euthanized by terminal cardiac puncture. Whole blood was allowed to clot at room temperature for 30 minutes, incubated at 37°C for 10 minutes, and then incubated at 4°C for 15 minutes before centrifugation (3,000 g, 20 minutes). Serum was stored at −80°C until analysis.

### ELISA measurements of soluble BAFF and LBP

Serum B-cell activation factor (BAFF) and lipopolysaccharide-binding protein (LBP) were quantified using the Quantikine Mouse BAFF/BLyS/TNFSF13B ELISA kit (R&D Systems) and the Mouse LBP ELISA kit (Abcam), respectively, following the manufacturers’ instructions.

### Colon epithelial cell isolation

Colon epithelial cells were isolated using a dithiothreitol (DTT)/EDTA-based epithelial stripping approach. Colons were washed in HBSS without Ca^++^ and Mg^++^ supplemented with EDTA (2 mM) and HEPES (25 mM). Tissue was incubated at 37°C for 15 minutes in stripping buffer (HBSS without Ca^++^/Mg^++^ supplemented with HEPES (15mM), EDTA (5mM) and DTT (1mM)) and vortexed to release epithelial cells. The epithelial fraction was filtered (100 μm), washed in PBS, and lysed in RLT buffer (QIAGEN) for RNA extraction.

### Tissue embedding, immunofluorescence staining and imaging

Colon tissue and brain hemisphere were embedded in OCT embedding medium (Scigen Scientific) and stored at −80°C. Colon cryosections (10µm) were prepared at the IRCM Histology Core Facility, fixed in cold acetone (−20°C), air dried, and stored at −80°C. Cryosections (14 μm) of brain hemispheres were prepared at the *Institut de Recherche en Immunologie et en Cancérologie* (IRIC) Histology Core Facility and processed identically.

Immunofluorescence staining was performed at the Centre de Recherche du Centre Hospitalier de l’Université de Montréal (CRCHUM) Molecular Pathology Core Facility using a Discovery Ultra automated stainer (Ventana/Roche). Sections were blocked in PBS containing 1% bovine serum albumin (BSA) for 30 minutes at room temperature, incubated with primary antibodies followed by species-appropriate secondary antibodies for 2 hours each at room temperature. Nuclei were counterstained with DAPI (1:3,000) for 10 minutes. Sudan Black (0.1% in 70% ethanol) was applied to reduce autofluorescence, and sections were mounted with Fluoromount (Sigma).

Slides were scanned using an Aperio Verso 200 scanner (Leica Biosystems) with a 20×/0.8 NA objective at a resolution of 0.275 μm per pixel. Image visualization and analysis were performed blinded to group allocation using Aperio ImageScope software (Leica Biosystems).

Primary antibodies used included rabbit anti-mouse zonula occludens-1 (ZO-1) and ionized calcium-binding adaptor molecule 1 (IBA1) and an anti-glial fibrillary acidic protein (GFAP) monoclonal antibody; Alexa Fluor-conjugated goat anti-rabbit secondary antibodies were used as indicated. Antibody details are provided in Table S4B.

### Cell culture and stimulation with GMEVs

#### THP-1-derived human macrophages

THP-1 cells were maintained in complete RPMI 1640 supplemented with 10% heat-inactivated fetal bovine serum (FBS), 1% penicillin-streptomycin and 14.3 μM 2-mercaptoethanol. For differentiation, THP-1 cells were seeded at 5 × 10^5^ cells per well (12-well plates) and treated with phorbol 12-myristate 13-acetate (PMA; 100nM) for 48h, washed and rested in fresh complete medium before stimulation. Macrophages were stimulated with GMEVs (1 μg/mL) for 5 h (RNA) or 16 h (secreted proteins). For intracellular cytokine staining, cells were treated with BD GolgiPlug protein transport inhibitor (containing brefeldin A) during the final 5h of stimulation.

#### Human intestinal organoids and epithelial monolayers

Human intestinal organoids (HIOs) were generated from human induced pluripotent stem cells (iPSCs) derived from fibroblasts from a healthy female donor (generously provided by H. Malech, NIH) using the STEMdiff Intestinal Organoid Kit (STEMCELL Technologies) following the manufacturer’s instructions. iPSCs were maintained on Matrigel in mTeSR1 medium and differentiated into intestinal lineage using the STEMdiff Intestinal Organoid Kit following the manufacturer’s protocol, including definitive endoderm induction and subsequent mid-hindgut patterning. Mid-hindgut spheroids were embedded in Matrigel domes and matured in Intestinal Organoid Growth Medium, with medium changes every 2-3 days and weekly passaging. For stimulation experiments, HIOs were dissociated into epithelial monolayers, plated onto Matrigel-coated 24-well plates and treated with GMEVs (1 µg/mL) for 5 h (RNA) or 16 h (secreted proteins).

### Transepithelial electrical resistance

Caco-2 cells were seeded at 1.5×10^5^ cells/cm^2^ on Transwell inserts (0.4 μm pore; 12-well format; Sarstedt) and maintained for ≥3 weeks until transepithelial electrical resistance (TEER) stabilized. Cells were cultured in Eagle’s Modified Essential Medium (EMEM) supplemented with 20% heat-inactivated FBS and penicillin-streptomycin. Baseline TEER was recorded before treatment. Based on preliminary dose–response optimization experiments, cells were treated with GMEVs (3µg/mL), and TEER measured 24h later using an EVOM2 voltohmmeter with STX2 electrodes. Values reflect the mean of 3 measurements per insert.

### RNA Extraction and quantitative PCR

Total RNA was extracted from colon epithelial cells, HIOs, iPSC-derived microglia, and THP-1-derived macrophages using the RNeasy Plus Mini Kit (QIAGEN) according to the manufacturer’s instructions. RNA concentration and purity were assessed using a NanoDrop 2000 Spectrophotometer (Thermo Fisher Scientific). Complementary DNA was synthesized from 1μg total RNA using the iScript Reverse Transcription SuperMix for RT-qPCR (BioRad). Quantitative PCR was performed using SsoAdvanced Universal SYBR Green Supermix (BioRad) on a StepOnePlus Real-Time PCR System (Applied Biosystems). Relative gene expression was calculated using the 2^(−ΔΔCt)^ method with GAPDH as the reference gene. Primer sequences are listed in Table S3.

### Differentiation of induced pluripotent stem cell-derived microglia

Human iPSCs were generated from a healthy female donor and maintained using standard procedures as described^73^. Microglia were differentiated from iPSCs using a two-step protocol adapted from established methods^74^. Briefly, iPSCs were differentiated into iPSC-derived hematopoietic progenitor cells (iHPCs) using the STEMdiff Hematopoietic Kit (STEMCELL Technologies), with minor modifications. On day −1, iPSCs were dissociated using Gentle Cell Dissociation Reagent (STEMCELL Technologies) and seeded onto Matrigel-coated 6-well plates in mTeSR Plus or Essential 8 medium supplemented with Y-27632 (10 μM, Selleckchem) at a density designed to yield colonies of fewer than 100 cells per cm² by day 0. On day 0, medium (1 mL) was replaced with STEMdiff Hematopoietic Medium A (2 mL per well). On day 2, half of the medium (1 mL) was replaced with fresh Medium A. On day 3, cultures were switched entirely to STEMdiff Hematopoietic Medium B (2mL per well). On days 5 and 7, half of the supernatant (1 mL) was replaced with fresh Medium B. On day 9, an additional 1 mL of fresh Medium B was added. On day 10, non-adherent iHPCs present in the supernatant were collected, centrifuged at 300 g for 5 minutes, and either cryopreserved (Bambanker, Fujifilm Wako Chemicals) or used for microglial differentiation. This harvesting procedure was repeated on days 12 and 14. For microglial differentiation, iHPCs were resuspended in microglia differentiation medium (as previously described) at a density of 5×10^4^ cells/mL and plated onto Matrigel-coated 6-well plates (2 mL per well), defining day 0 of differentiation. Cultures were supplemented every other day with 1mL of fresh differentiation medium from day 0 to day 10. On day 12, supernatants were collected, cells were pelleted by centrifugation (300 g, 5 minutes), resuspended in fresh differentiation medium, and returned to culture. This procedure was repeated on day 24. Cells were considered mature by day 28 and were maintained with media changes every other day until use. All cultures were maintained at 37°C in a humidified atmosphere containing 5% CO2. For downstream experiments, cells were detached using PBS containing 2mM EDTA and replated at the desired density.

### Bulk RNA sequencing of GMEV-treated iMGLs

Induced pluripotent stem cell-derived microglia (iMGLs) were treated with GMEVs (1 μg/mL) from LC donors (n = 10) or PC donors (n = 4) for 16 h. RNA (≤100ng) was submitted for ribodepletion library preparation and sequencing (target ∼50 million reads per sample; IRCM Molecular Biology platform). Read quality was assessed using FastQC (v0.12.1). Reads were aligned to GRCh38 using STAR (v2.7.11b), gene counts were generated using featureCounts v2.0.6; GRCh38 release 110), and differential expression was performed using the DESeq2. Heatmaps were generated using z-scored normalized counts. Functional enrichment analysis was performed using gprofiler2.

### Cytokine and chemokine quantification in cell culture supernatants

Secreted cytokines/chemokines were quantified using MSD U-PLEX panels. For THP-1 macrophages and HIOs, analytes included granulocyte-macrophage colony stimulating factor (GM-CSF), granulocyte-colony stimulating factor (G-CSF), interferon gamma (IFNγ), IL-10, IL-1β, IL-6, IL-8, CXCL10, CCL2, and tumor necrosis factor (TNF). For iMGL supernatants, the panel included B-cell activation factor (BAFF), IL-1β, IL-6, IL-8, TNF, s100A12, matrix metalloproteinase 9 (MMP9), CCL2 and CXCL1, with R-PLEX detection of C1q. Plates were read on a MESO QuickPlex SQ 120 and analyzed using MSD Discovery Workbench 4.0.

### Flow cytometry of THP-1-derived macrophages

Cells were stained with LIVE/DEAD Fixable Aqua (Thermo Fisher Scientific), blocked with human Fc block (BD Biosciences) with 20% heat inactivated FBS and 50 μg mouse and/or rat IgG, and stained for surface BAFF, followed by fixation/permeabilization (Cytofix/Cytoperm, BD Biosciences) and intracellular staining for IL-1ꞵ, IL-6 and TNF. Data were acquired on a BD LSRFortessa and analyzed in FlowJo (v10.8.1). Antibody details are provided in Table S4A.

### Statistical analyses

Group comparisons were performed using one-way ANOVA with Tukey’s post hoc test for approximately normally distributed data, or Kruskal-Wallis tests with Dunn’s post hoc correction otherwise. For two-group comparisons, Wilcoxon rank-sum (two-sided) tests were used unless stated otherwise. Correlations were assessed using Pearson’s or Spearman’s tests, as appropriate. Categorical variables were analyzed using Fisher’s exact test. Unless otherwise indicated, data are shown as medians with interquartile ranges. All tests were two-sided and *P* < 0.05 was considered statistically significant. Analyses were performed using GraphPad Prism v10.2.0.

## Acknowledgements

E.L.F is supported by a Tier 2 Canada Research Chair in Role of the Microbiome in Inborn Errors of Immunity and Post-Infectious Conditions, the Canadian Institutes of Health Research (CIHR), the *Fonds de Recherche du Québec (FRQ)*. K.D.L. is supported by the IRCM Foundation. M.A. is supported by CIHR. A.D. is supported by the *Fonds de Recherche du Québec (FRQ)*. I.B. is supported by the IRCM Foundation. The work was supported by CIHR (PJT-191724), the Canada Research Chairs Program, the FRQ Clinical Research Scholars - Junior 1 Establishment Funds for Young Investigators, the John R. Evans Leaders Fund from the Canadian Foundation for Innovation (CFI), the J-Louis Lévesque Foundation Research Chair, the Mirella and Lino Saputo Foundation, and the IRCM Foundation.

We thank members of the IRCM animal facility (Mariane Canuel, Eve-Marie Charbonneau, Manon Laprise, and Jo-Anny Bisson), the IRCM Flow Cytometry Platform (Éric Massicotte and Julie Lord), and the McGill Genome Center, Center for Microbiome Research, for technical support. We are grateful to Kathy McCoy for providing germ-free mice. We thank Christian Charbonneau, Dr. Marianne Isaac and Melina Narlis of the IRIC Microscopy and Histology Core Facilities for guidance and for performing sagittal brain cryosections and H&E staining. We also thank Véronique Barrès and Liliane Meunier of the CRCHUM Molecular Pathology Core Facility for immunolabeling, slide scanning, and assistance with paraffin and OCT processing of colon and brain, and Anabelle Bouchard-Bourque of the IRCM Histology Core Facility for additional paraffin and OCT sectioning. We acknowledge Dominic Filion and Mattew Duguay of the IRCM Imaging Platform for imaging support. We thank Dr. Mieczyslaw Marcinkiewicz, and Dennis A. Drewnik for advice and assistance with brain immunofluorescence interpretation, fixation and embedding. We also thank the Center for Applied Nanomedicine at the Research Institute of the McGill University Hospital Center for assistance with nanoparticle tracking analysis, and Dr. S. Kelly Sears and Dr. Jeannie Mui from the Facility for Electron Microscopy Research at McGill University for their assistance with electron microscopy.

## Author contributions

M.A. and E.L.F. conceived the study and designed the research. M.A., K.D-L., I.B., A.V., A.D., E.D, L.P., V.E.P., F.B., J.S., and A.A. performed experiments. M.A., K.D-L., I.B., P.C., J.P. and E.L.F. analyzed the data. M.A., P.C., and E.L.F. wrote the initial manuscript draft, and all authors contributed to manuscript revision. E.L.F. secured funding for the study. E.L.F. supervised the research.

## Data availability

All raw sequencing data has been deposited in SRA under BioProject PRJNA1236664 (submissions: SUB15113758 and SUB15164815) and will be released upon publication. Any additional data supporting the findings of this study are available from the corresponding author upon reasonable request.

## Code availability

Custom code used for microbiota processing, statistical analyses, and figure generation will be deposited on GitHub and made publicly available upon publication. Scripts are available from the corresponding author upon reasonable request prior to release.

## References

1. Kuang, S.E., S.; Clarke, J.; Zakaria, D.; Demers, A.; Aziz, S. (2023). Experiences of Canadians with long-term symptoms following COVID-19. Insights on Canadian Society. Published online December 8, 2023.

2. Ely, E.W., Brown, L.M., Fineberg, H.V., National Academies of Sciences, E., and Medicine Committee on Examining the Working Definition for Long, C. (2024). Long Covid Defined. N Engl J Med 391, 1746–1753. 10.1056/NEJMsb2408466.

3. COVID-19, W.H.O.W.c.c.d.w.g.o.p., and condition (2021). A clinical case definition of post COVID-19 condition by a Delphi consensus.

4. Davis, H.E., Assaf, G.S., McCorkell, L., Wei, H., Low, R.J., Re’em, Y., Redfield, S., Austin, J.P., and Akrami, A. (2021). Characterizing long COVID in an international cohort: 7 months of symptoms and their impact. EClinicalMedicine 38, 101019. 10.1016/j.eclinm.2021.101019.

5. Lopez-Leon, S., Wegman-Ostrosky, T., Perelman, C., Sepulveda, R., Rebolledo, P.A., Cuapio, A., and Villapol, S. (2021). More than 50 long-term effects of COVID-19: a systematic review and meta-analysis. Sci Rep 11, 16144. 10.1038/s41598-021-95565-8.

6. Davis, H.E., McCorkell, L., Vogel, J.M., and Topol, E.J. (2023). Long COVID: major findings, mechanisms and recommendations. Nat Rev Microbiol 21, 133–146. 10.1038/s41579-022-00846-2.

7. Al-Aly, Z., Xie, Y., and Bowe, B. (2021). High-dimensional characterization of post-acute sequelae of COVID-19. Nature 594, 259–264. 10.1038/s41586-021-03553-9.

8. Al-Aly, Z., and Topol, E. (2024). Solving the puzzle of Long Covid. Science 383, 830–832. 10.1126/science.adl0867.

9. Cai, M., Xie, Y., Topol, E.J., and Al-Aly, Z. (2024). Three-year outcomes of post-acute sequelae of COVID-19. Nat Med 30, 1564–1573. 10.1038/s41591-024-02987-8.

10. Klein, J., Wood, J., Jaycox, J.R., Dhodapkar, R.M., Lu, P., Gehlhausen, J.R., Tabachnikova, A., Greene, K., Tabacof, L., Malik, A.A., et al. (2023). Distinguishing features of long COVID identified through immune profiling. Nature 623, 139–148. 10.1038/s41586-023-06651-y.

11. Yin, K., Peluso, M.J., Luo, X., Thomas, R., Shin, M.G., Neidleman, J., Andrew, A., Young, K.C., Ma, T., Hoh, R., et al. (2024). Long COVID manifests with T cell dysregulation, inflammation and an uncoordinated adaptive immune response to SARS-CoV-2. Nat Immunol 25, 218–225. 10.1038/s41590-023-01724-6.

12. Phetsouphanh, C., Darley, D.R., Wilson, D.B., Howe, A., Munier, C.M.L., Patel, S.K., Juno, J.A., Burrell, L.M., Kent, S.J., Dore, G.J., et al. (2022). Immunological dysfunction persists for 8 months following initial mild-to-moderate SARS-CoV-2 infection. Nat Immunol 23, 210–216. 10.1038/s41590-021-01113-x.

13. Woodruff, M.C., Bonham, K.S., Anam, F.A., Walker, T.A., Faliti, C.E., Ishii, Y., Kaminski, C.Y., Ruunstrom, M.C., Cooper, K.R., Truong, A.D., et al. (2023). Chronic inflammation, neutrophil activity, and autoreactivity splits long COVID. Nat Commun 14, 4201. 10.1038/s41467-023-40012-7.

14. Talla, A., Vasaikar, S.V., Szeto, G.L., Lemos, M.P., Czartoski, J.L., MacMillan, H., Moodie, Z., Cohen, K.W., Fleming, L.B., Thomson, Z., et al. (2023). Persistent serum protein signatures define an inflammatory subcategory of long COVID. Nat Commun 14, 3417. 10.1038/s41467-023-38682-4.

15. Pretorius, E., Vlok, M., Venter, C., Bezuidenhout, J.A., Laubscher, G.J., Steenkamp, J., and Kell, D.B. (2021). Persistent clotting protein pathology in Long COVID/Post-Acute Sequelae of COVID-19 (PASC) is accompanied by increased levels of antiplasmin. Cardiovasc Diabetol 20, 172. 10.1186/s12933-021-01359-7.

16. Braga, J., Lepra, M., Kish, S.J., Rusjan, P.M., Nasser, Z., Verhoeff, N., Vasdev, N., Bagby, M., Boileau, I., Husain, M.I., et al. (2023). Neuroinflammation After COVID-19 With Persistent Depressive and Cognitive Symptoms. JAMA Psychiatry 80, 787–795. 10.1001/jamapsychiatry.2023.1321.

17. Soung, A.L., Vanderheiden, A., Nordvig, A.S., Sissoko, C.A., Canoll, P., Mariani, M.B., Jiang, X., Bricker, T., Rosoklija, G.B., Arango, V., et al. (2022). COVID-19 induces CNS cytokine expression and loss of hippocampal neurogenesis. Brain 145, 4193–4201. 10.1093/brain/awac270.

18. Chang, R., Yen-Ting Chen, T., Wang, S.I., Hung, Y.M., Chen, H.Y., and Wei, C.J. (2023). Risk of autoimmune diseases in patients with COVID-19: A retrospective cohort study. EClinicalMedicine 56, 101783. 10.1016/j.eclinm.2022.101783.

19. Son, K., Jamil, R., Chowdhury, A., Mukherjee, M., Venegas, C., Miyasaki, K., Zhang, K., Patel, Z., Salter, B., Yuen, A.C.Y., et al. (2023). Circulating anti-nuclear autoantibodies in COVID-19 survivors predict long COVID symptoms. Eur Respir J 61. 10.1183/13993003.00970-2022.

20. Nayyerabadi, M., Fourcade, L., Joshi, S.A., Chandrasekaran, P., Chakravarti, A., Masse, C., Paul, M.L., Houle, J., Boubekeur, A.M., DuSablon, C., et al. (2023). Vaccination after developing long COVID: Impact on clinical presentation, viral persistence, and immune responses. Int J Infect Dis 136, 136–145. 10.1016/j.ijid.2023.09.006.

21. Swank, Z., Senussi, Y., Manickas-Hill, Z., Yu, X.G., Li, J.Z., Alter, G., and Walt, D.R. (2023). Persistent Circulating Severe Acute Respiratory Syndrome Coronavirus 2 Spike Is Associated With Post-acute Coronavirus Disease 2019 Sequelae. Clin Infect Dis 76, e487–e490. 10.1093/cid/ciac722.

22. Natarajan, A., Zlitni, S., Brooks, E.F., Vance, S.E., Dahlen, A., Hedlin, H., Park, R.M., Han, A., Schmidtke, D.T., Verma, R., et al. (2022). Gastrointestinal symptoms and fecal shedding of SARS-CoV-2 RNA suggest prolonged gastrointestinal infection. Med 3, 371–387 e379. 10.1016/j.medj.2022.04.001.

23. Stein, S.R., Ramelli, S.C., Grazioli, A., Chung, J.Y., Singh, M., Yinda, C.K., Winkler, C.W., Sun, J., Dickey, J.M., Ylaya, K., et al. (2022). SARS-CoV-2 infection and persistence in the human body and brain at autopsy. Nature 612, 758–763. 10.1038/s41586-022-05542-y.

24. Zollner, A., Koch, R., Jukic, A., Pfister, A., Meyer, M., Rossler, A., Kimpel, J., Adolph, T.E., and Tilg, H. (2022). Postacute COVID-19 is Characterized by Gut Viral Antigen Persistence in Inflammatory Bowel Diseases. Gastroenterology 163, 495–506 e498. 10.1053/j.gastro.2022.04.037.

25. Yeoh, Y.K., Zuo, T., Lui, G.C., Zhang, F., Liu, Q., Li, A.Y., Chung, A.C., Cheung, C.P., Tso, E.Y., Fung, K.S., et al. (2021). Gut microbiota composition reflects disease severity and dysfunctional immune responses in patients with COVID-19. Gut 70, 698–706. 10.1136/gutjnl-2020-323020.

26. Liu, Q., Mak, J.W.Y., Su, Q., Yeoh, Y.K., Lui, G.C., Ng, S.S.S., Zhang, F., Li, A.Y.L., Lu, W., Hui, D.S., et al. (2022). Gut microbiota dynamics in a prospective cohort of patients with post-acute COVID-19 syndrome. Gut 71, 544–552. 10.1136/gutjnl-2021-325989.

27. Honda, K., and Littman, D.R. (2016). The microbiota in adaptive immune homeostasis and disease. Nature 535, 75–84. 10.1038/nature18848.

28. Malard, F., Dore, J., Gaugler, B., and Mohty, M. (2021). Introduction to host microbiome symbiosis in health and disease. Mucosal Immunol 14, 547–554. 10.1038/s41385-020-00365-4.

29. Di Vincenzo, F., Del Gaudio, A., Petito, V., Lopetuso, L.R., and Scaldaferri, F. (2024). Gut microbiota, intestinal permeability, and systemic inflammation: a narrative review. Internal and Emergency Medicine 19, 275–293. 10.1007/s11739-023-03374-w.

30. Giron, L.B., Dweep, H., Yin, X., Wang, H., Damra, M., Goldman, A.R., Gorman, N., Palmer, C.S., Tang, H.Y., Shaikh, M.W., et al. (2021). Plasma Markers of Disrupted Gut Permeability in Severe COVID-19 Patients. Front Immunol 12, 686240. 10.3389/fimmu.2021.686240.

31. Prasad, R., Patton, M.J., Floyd, J.L., Fortmann, S., DuPont, M., Harbour, A., Wright, J., Lamendella, R., Stevens, B.R., Oudit, G.Y., and Grant, M.B. (2022). Plasma Microbiome in COVID-19 Subjects: An Indicator of Gut Barrier Defects and Dysbiosis. Int J Mol Sci 23. 10.3390/ijms23169141.

32. Giron, L.B., Peluso, M.J., Ding, J., Kenny, G., Zilberstein, N.F., Koshy, J., Hong, K.Y., Rasmussen, H., Miller, G.E., Bishehsari, F., et al. (2022). Markers of fungal translocation are elevated during post-acute sequelae of SARS-CoV-2 and induce NF-kappaB signaling. JCI Insight 7. 10.1172/jci.insight.160989.

33. Morais, L.H., Schreiber, H.L.t., and Mazmanian, S.K. (2021). The gut microbiota-brain axis in behaviour and brain disorders. Nat Rev Microbiol 19, 241–255. 10.1038/s41579-020-00460-0.

34. Ancuta, P., Kamat, A., Kunstman, K.J., Kim, E.Y., Autissier, P., Wurcel, A., Zaman, T., Stone, D., Mefford, M., Morgello, S., et al. (2008). Microbial translocation is associated with increased monocyte activation and dementia in AIDS patients. PLoS One 3, e2516. 10.1371/journal.pone.0002516.

35. Chen, X., Wei, J., Zhang, Y., Zhang, Y., and Zhang, T. (2024). Crosstalk between gut microbiome and neuroinflammation in pathogenesis of HIV-associated neurocognitive disorder. J Neurol Sci 457, 122889. 10.1016/j.jns.2024.122889.

36. Leclerc, L., Poudrier, J., Power, C., Lam, G.Y., and Falcone, E.L. (2025). Intestinal barrier compromise, viral persistence, and immune dysregulation converge on neurological sequelae in Long COVID. Front Aging Neurosci 17, 1744415. 10.3389/fnagi.2025.1744415.

37. Xie, J., Haesebrouck, F., Van Hoecke, L., and Vandenbroucke, R.E. (2023). Bacterial extracellular vesicles: an emerging avenue to tackle diseases. Trends Microbiol 31, 1206–1224. 10.1016/j.tim.2023.05.010.

38. Villard, A., Boursier, J., and Andriantsitohaina, R. (2021). Microbiota-derived extracellular vesicles and metabolic syndrome. Acta Physiol (Oxf) 231, e13600. 10.1111/apha.13600.

39. Villard, A., Boursier, J., and Andriantsitohaina, R. (2021). Bacterial and eukaryotic extracellular vesicles and nonalcoholic fatty liver disease: new players in the gut-liver axis? Am J Physiol Gastrointest Liver Physiol 320, G485–G495. 10.1152/ajpgi.00362.2020.

40. Fizanne, L., Villard, A., Benabbou, N., Recoquillon, S., Soleti, R., Delage, E., Wertheimer, M., Vidal-Gomez, X., Oullier, T., Chaffron, S., et al. (2023). Faeces-derived extracellular vesicles participate in the onset of barrier dysfunction leading to liver diseases. J Extracell Vesicles 12, e12303. 10.1002/jev2.12303.

41. Fasano, A. (2011). Zonulin and its regulation of intestinal barrier function: the biological door to inflammation, autoimmunity, and cancer. Physiol Rev 91, 151–175. 10.1152/physrev.00003.2008.

42. Kraeuter, A.K., Guest, P.C., and Sarnyai, Z. (2018). The Y-Maze for Assessment of Spatial Working and Reference Memory in Mice. Methods in molecular biology 1916, 105–111.

43. Kitsios, G.D., Li, K., Blacka, S., Fitch, A., Jacobs, J., Naqvi, A., Zhang, B., Gentry, H., Murray, C., Wang, X., et al. (2025). Oral and gut microbiota relate to symptom subphenotypes in long COVID, independent of viral persistence. iScience 28, 113628. 10.1016/j.isci.2025.113628.

44. Mendes de Almeida, V., Engel, D.F., Ricci, M.F., Cruz, C.S., Lopes Í, S., Alves, D.A., d’ Auriol1, M., Magalhães, J., Machado, E.C., Rocha, V.M., et al. (2023). Gut microbiota from patients with COVID-19 cause alterations in mice that resemble post-COVID symptoms. Gut Microbes 15, 2249146. 10.1080/19490976.2023.2249146.

45. Forte, D., Pellegrino, R.M., Trabanelli, S., Tonetti, T., Ricci, F., Cenerenti, M., Comai, G., Tazzari, P., Lazzarotto, T., Buratta, S., et al. (2023). Circulating extracellular particles from severe COVID-19 patients show altered profiling and innate lymphoid cell-modulating ability. Front Immunol 14, 1085610. 10.3389/fimmu.2023.1085610.

46. Nair, S., Nova-Lamperti, E., Labarca, G., Kulasinghe, A., Short, K.R., Carrión, F., and Salomon, C. (2023). Genomic communication via circulating extracellular vesicles and long-term health consequences of COVID-19. J Transl Med 21, 709. 10.1186/s12967-023-04552-2.

47. Jones, E.J., Booth, C., Fonseca, S., Parker, A., Cross, K., Miquel-Clopes, A., Hautefort, I., Mayer, U., Wileman, T., Stentz, R., and Carding, S.R. (2020). The Uptake, Trafficking, and Biodistribution of Bacteroides thetaiotaomicron Generated Outer Membrane Vesicles. Front Microbiol 11, 57. 10.3389/fmicb.2020.00057.

48. Doyon-Laliberte, K., Aranguren, M., Byrns, M., Chagnon-Choquet, J., Paniconi, M., Routy, J.P., Tremblay, C., Quintal, M.C., Brassard, N., Kaufmann, D.E., et al. (2022). Excess BAFF Alters NR4As Expression Levels and Breg Function of Human Precursor-like Marginal Zone B-Cells in the Context of HIV-1 Infection. Int J Mol Sci 23. 10.3390/ijms232315142.

49. Aranguren, M., Doyon-Laliberté, K., El-Far, M., Chartrand-Lefebvre, C., Routy, J.P., Barril, J.G., Trottier, B., Tremblay, C., Durand, M., Poudrier, J., and Roger, M. (2022). Subclinical Atherosclerosis Is Associated with Discrepancies in BAFF and APRIL Levels and Altered Breg Potential of Precursor-like Marginal Zone B-Cells in Long-Term HIV Treated Individuals. Vaccines (Basel) 11. 10.3390/vaccines11010081.

50. Becker-Merok, A., Nikolaisen, C., and Nossent, H.C. (2006). B-lymphocyte activating factor in systemic lupus erythematosus and rheumatoid arthritis in relation to autoantibody levels, disease measures and time. Lupus 15, 570–576. 10.1177/0961203306071871.

51. Chu, V.T., Enghard, P., Schürer, S., Steinhauser, G., Rudolph, B., Riemekasten, G., and Berek, C. (2009). Systemic activation of the immune system induces aberrant BAFF and APRIL expression in B cells in patients with systemic lupus erythematosus. Arthritis & Rheumatism 60, 2083–2093. 10.1002/art.24628.

52. Steri, M., Orrù, V., Idda, M.L., Pitzalis, M., Pala, M., Zara, I., Sidore, C., Faà, V., Floris, M., Deiana, M., et al. (2017). Overexpression of the Cytokine BAFF and Autoimmunity Risk. N Engl J Med 376, 1615–1626. 10.1056/NEJMoa1610528.

53. Lesley, R., Xu, Y., Kalled, S.L., Hess, D.M., Schwab, S.R., Shu, H.B., and Cyster, J.G. (2004). Reduced competitiveness of autoantigen-engaged B cells due to increased dependence on BAFF. Immunity 20, 441–453. 10.1016/s1074-7613(04)00079-2.

54. Doyon-Laliberté, K., Aranguren, M., Chagnon-Choquet, J., Batraville, L.A., Dagher, O., Richard, J., Paniconi, M., Routy, J.P., Tremblay, C., Quintal, M.C., et al. (2023). Excess BAFF May Impact HIV-1-Specific Antibodies and May Promote Polyclonal Responses Including Those from First-Line Marginal Zone B-Cell Populations. Curr Issues Mol Biol 46, 25–43. 10.3390/cimb46010003.

55. Brenchley, J.M., Price, D.A., Schacker, T.W., Asher, T.E., Silvestri, G., Rao, S., Kazzaz, Z., Bornstein, E., Lambotte, O., Altmann, D., et al. (2006). Microbial translocation is a cause of systemic immune activation in chronic HIV infection. Nat Med 12, 1365–1371. 10.1038/nm1511.

56. Isnard, S., Ramendra, R., Dupuy, F.P., Lin, J., Fombuena, B., Kokinov, N., Kema, I., Jenabian, M.-A., Lebouché, B., and Costiniuk, C.T. (2020). Plasma levels of C-type lectin REG3α and gut damage in people with human immunodeficiency virus. The Journal of infectious diseases 221, 110–121.

57. Steimle, A., Autenrieth, I.B., and Frick, J.-S. (2016). Structure and function: Lipid A modifications in commensals and pathogens. International Journal of Medical Microbiology 306, 290–301. 10.1016/j.ijmm.2016.03.001.

58. Behrouzi, A., Vaziri, F., Riazi Rad, F., Amanzadeh, A., Fateh, A., Moshiri, A., Khatami, S., and Siadat, S.D. (2018). Comparative study of pathogenic and non-pathogenic Escherichia coli outer membrane vesicles and prediction of host-interactions with TLR signaling pathways. BMC Res Notes 11, 539. 10.1186/s13104-018-3648-3.

59. Andrews, S. (2010). FastQC: a quality control tool for high throughput sequence data.

60. Franzosa, E.A., McIver, L.J., Rahnavard, G., Thompson, L.R., Schirmer, M., Weingart, G., Lipson, K.S., Knight, R., Caporaso, J.G., Segata, N., and Huttenhower, C. (2018). Species-level functional profiling of metagenomes and metatranscriptomes. Nat Methods 15, 962–968. 10.1038/s41592-018-0176-y.

61. Manghi, P., Blanco-Míguez, A., Manara, S., NabiNejad, A., Cumbo, F., Beghini, F., Armanini, F., Golzato, D., Huang, K.D., Thomas, A.M., et al. (2023). MetaPhlAn 4 profiling of unknown species-level genome bins improves the characterization of diet-associated microbiome changes in mice. Cell Reports 42, 112464. 10.1016/j.celrep.2023.112464.

62. McMurdie, P.J., and Holmes, S. (2013). phyloseq: an R package for reproducible interactive analysis and graphics of microbiome census data. PLoS One 8, e61217. 10.1371/journal.pone.0061217.

63. Whitmore, N. (2022). R for Conservation and Development Projects A Primer for Practitioners (Chapman and Hall / CRC Press).

64. Robinson, M.D., McCarthy, D.J., and Smyth, G.K. (2010). edgeR: a Bioconductor package for differential expression analysis of digital gene expression data. Bioinformatics 26, 139–140. 10.1093/bioinformatics/btp616.

65. Cao, Y., Dong, Q., Wang, D., Zhang, P., Liu, Y., and Niu, C. (2022). microbiomeMarker: an R/Bioconductor package for microbiome marker identification and visualization. Bioinformatics 38, 4027–4029. 10.1093/bioinformatics/btac438.

66. Bolger, A.M., Lohse, M., and Usadel, B. (2014). Trimmomatic: a flexible trimmer for Illumina sequence data. Bioinformatics 30, 2114–2120. 10.1093/bioinformatics/btu170.

67. Masella, A.P., Bartram, A.K., Truszkowski, J.M., Brown, D.G., and Neufeld, J.D. (2012). PANDAseq: paired-end assembler for illumina sequences. BMC Bioinformatics 13, 31. 10.1186/1471-2105-13-31.

68. Mlaga, K.D., Mathieu, A., Beauparlant, C.J., Ott, A., Khodr, A., Perin, O., and Droit, A. (2021). HCK and ABAA: A Newly Designed Pipeline to Improve Fungi Metabarcoding Analysis. Front Microbiol 12, 640693. 10.3389/fmicb.2021.640693.

69. Fu, L., Niu, B., Zhu, Z., Wu, S., and Li, W. (2012). CD-HIT: accelerated for clustering the next-generation sequencing data. Bioinformatics 28, 3150–3152. 10.1093/bioinformatics/bts565.

70. Lu, J., and Salzberg, S.L. (2020). Ultrafast and accurate 16S rRNA microbial community analysis using Kraken 2. Microbiome 8, 124. 10.1186/s40168-020-00900-2.

71. Dixon, P. (2003). VEGAN, a package of R functions for community ecology. Journal of vegetation science 14, 927–930. 10.1111/j.1654-1103.2003.tb02228.x.

72. Seibenhener, M.L., and Wooten, M.C. (2015). Use of the Open Field Maze to measure locomotor and anxiety-like behavior in mice. J Vis Exp, e52434. 10.3791/52434.

73. Chen, C.X., Abdian, N., Maussion, G., Thomas, R.A., Demirova, I., Cai, E., Tabatabaei, M., Beitel, L.K., Karamchandani, J., Fon, E.A., and Durcan, T.M. (2021). A Multistep Workflow to Evaluate Newly Generated iPSCs and Their Ability to Generate Different Cell Types. Methods Protoc 4 10.3390/mps4030050.

74. Dorion, M.-F., Casas, D., Shlaifer, I., Yaqubi, M., Fleming, P., Karpilovsky, N., Chen, C.X.Q., Nicouleau, M., Piscopo, V.E.C., MacDougall, E.J., et al. (2024). An adapted protocol to derive microglia from stem cells and its application in the study of CSF1R-related disorders. Molecular Neurodegeneration 19, 31. 10.1186/s13024-024-00723-x.

