## Supplemental Figures and Tables for "Microbiota-derived extracellular vesicles link intestinal dysbiosis to neuroimmune activation in long COVID"

Fig.S1

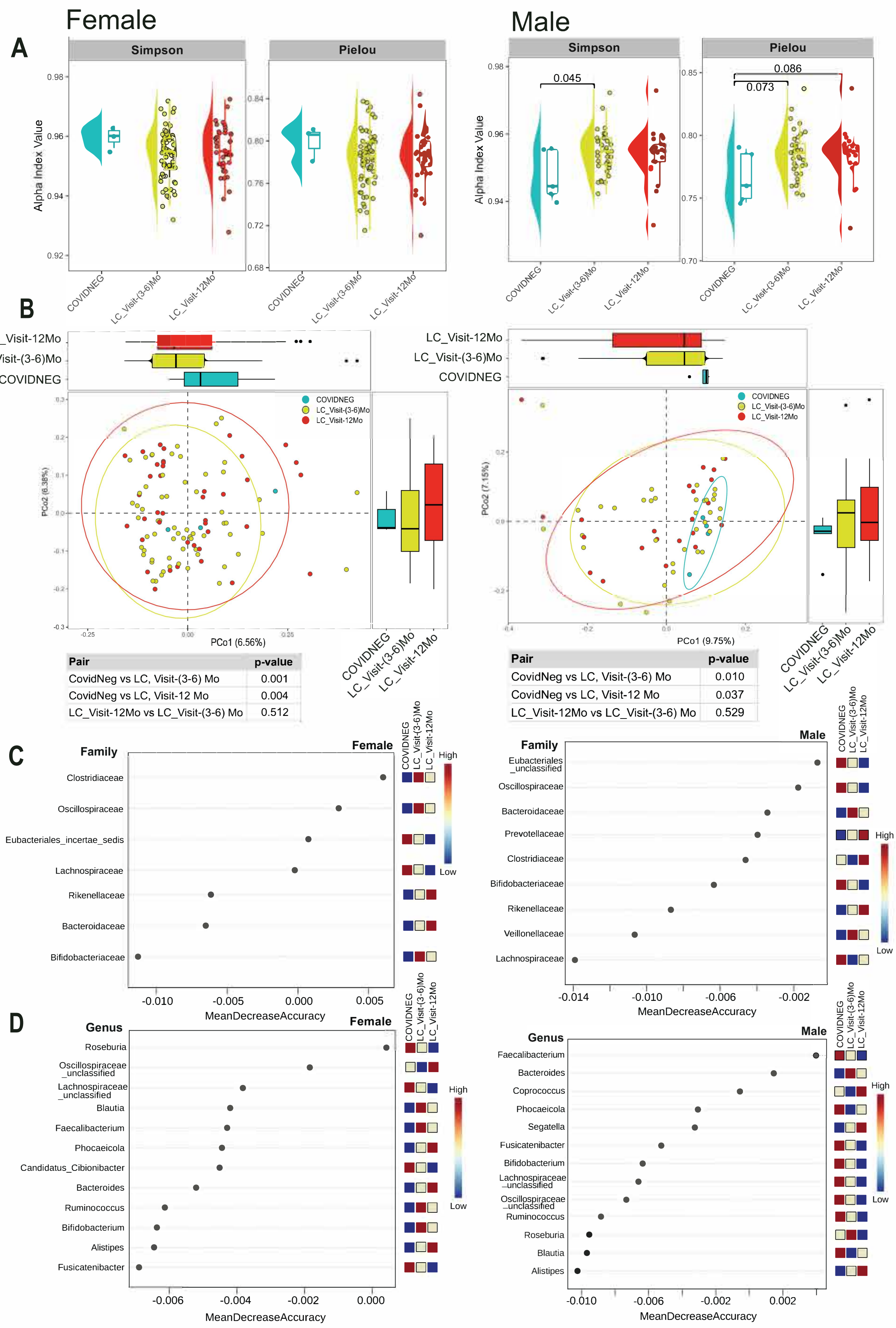


| Pair | p-value |
|---|---|
| CovidNeg vs LC, Visit-(3-6) Mo | 0.001 |
| CovidNeg vs LC, Visit-12 Mo | 0.004 |
| LC_Visit-12Mo vs LC_Visit-(3-6) Mo | 0.512 |

| Pair | p-value |
|---|---|
| CovidNeg vs LC, Visit-(3-6) Mo | 0.010 |
| CovidNeg vs LC, Visit-12 Mo | 0.037 |
| LC_Visit-12Mo vs LC_Visit-(3-6) Mo | 0.529 |

Fig.S2

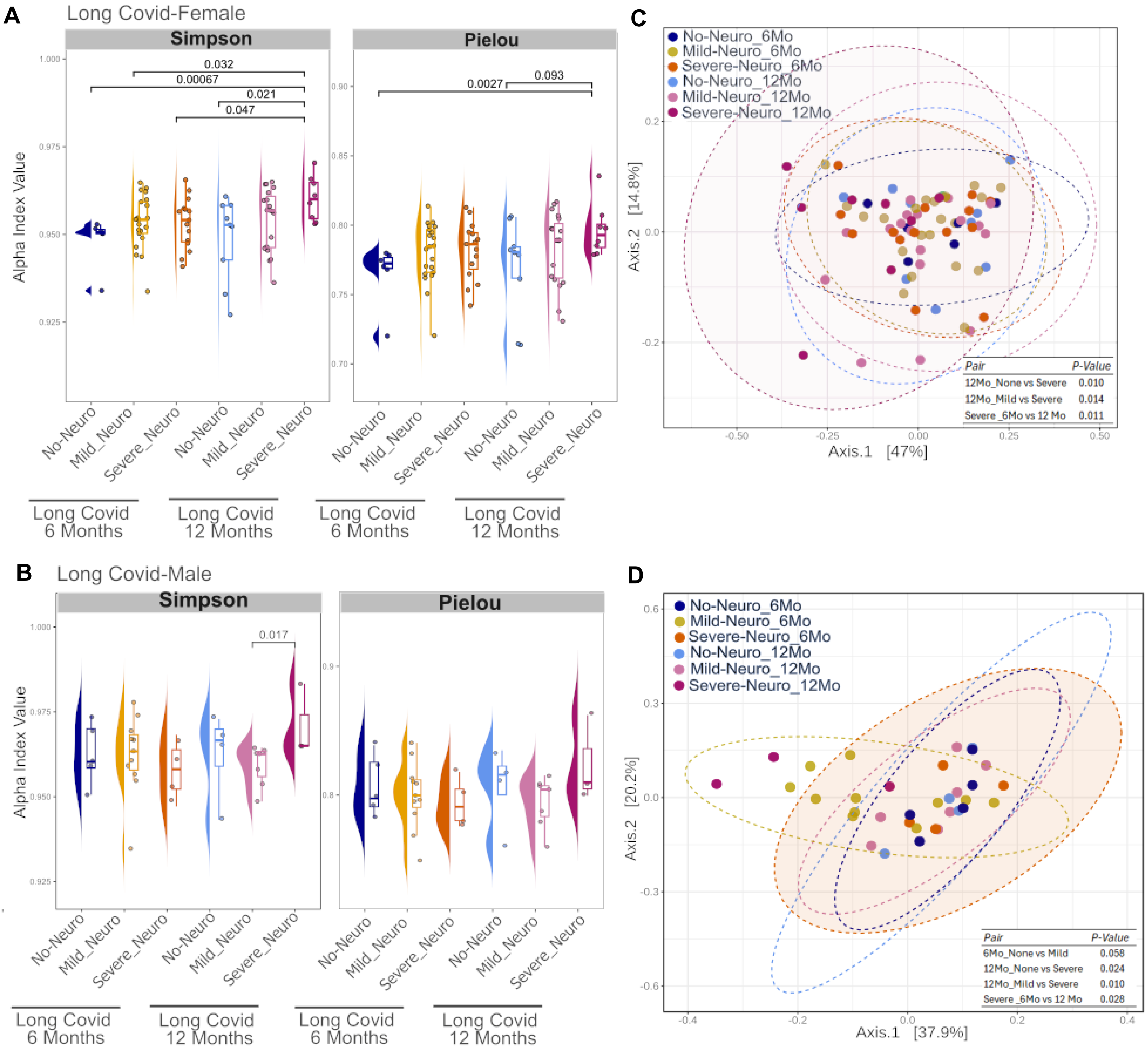
A
Long Covid-Female
Simpson
Pielou
Alpha Index Value
0.032
0.00067
0.021
0.047
0.0027
0.093
No-Neuro
Mild_Neuro
Severe_Neuro
Long Covid 6 Months
Long Covid 12 Months
B
Long Covid-Male
0.017
C
No-Neuro_6Mo
Mild-Neuro_6Mo
Severe-Neuro_6Mo
No-Neuro_12Mo
Mild-Neuro_12Mo
Severe-Neuro_12Mo
Axis.1 [47%]
Axis.2 [14.8%]
Pair P-Value
12Mo_None vs Severe 0.010
12Mo_Mild vs Severe 0.014
Severe _6Mo vs 12 Mo 0.011
D
Axis.1 [37.9%]
Axis.2 [20.2%]
Pair P-Value
6Mo_None vs Mild 0.058
12Mo_None vs Severe 0.024
12Mo_Mild vs Severe 0.010
Severe _6Mo vs 12 Mo 0.028

**Fig.S3**

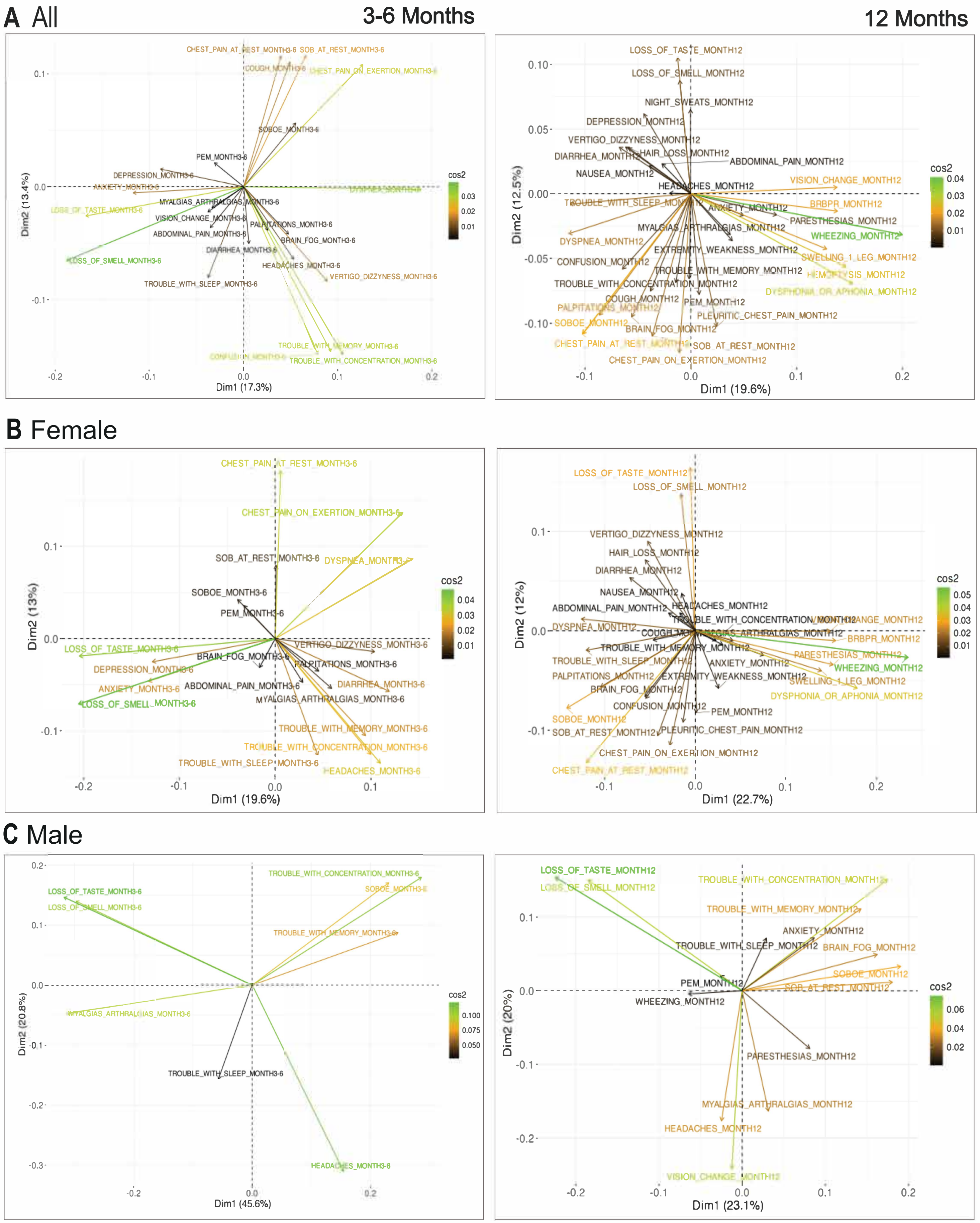
A All
3-6 Months
12 Months
CHEST_PAIN_AT_REST_MONTH3-6
SOB_AT_REST_MONTH3-6
COUGH_MONTH3-6
CHEST_PAIN_ON_EXERTION_MONTH3-6
SOBOE_MONTH3-6
PEM_MONTH3-6
DEPRESSION_MONTH3-6
ANXIETY_MONTH3-6
DYSPNEA_MONTH3-6
LOSS_OF_TASTE_MONTH3-6
MYALGIAS_ARTHRALGIAS_MONTH3-6
VISION_CHANGE_MONTH3-6
PALPITATIONS_MONTH3-6
ABDOMINAL_PAIN_MONTH3-6
BRAIN_FOG_MONTH3-6
DIARRHEA_MONTH3-6
LOSS_OF_SMELL_MONTH3-6
HEADACHES_MONTH3-6
VERTIGO_DIZZYNESS_MONTH3-6
TROUBLE_WITH_SLEEP_MONTH3-6
TROUBLE_WITH_MEMORY_MONTH3-6
CONFUSION_MONTH3-6
TROUBLE_WITH_CONCENTRATION_MONTH3-6
Dim1 (17.3%)
Dim2 (13.4%)
cos2
LOSS_OF_TASTE_MONTH12
LOSS_OF_SMELL_MONTH12
NIGHT_SWEATS_MONTH12
DEPRESSION_MONTH12
VERTIGO_DIZZYNESS_MONTH12
DIARRHEA_MONTH12
HAIR_LOSS_MONTH12
ABDOMINAL_PAIN_MONTH12
NAUSEA_MONTH12
HEADACHES_MONTH12
VISION_CHANGE_MONTH12
TROUBLE_WITH_SLEEP_MONTH12
ANXIETY_MONTH12
BRBPR_MONTH12
PARESTHESIAS_MONTH12
MYALGIAS_ARTHRALGIAS_MONTH12
WHEEZING_MONTH12
DYSPNEA_MONTH12
EXTREMITY_WEAKNESS_MONTH12
SWELLING_1_LEG_MONTH12
CONFUSION_MONTH12
TROUBLE_WITH_MEMORY_MONTH12
HEMOPTYSIS_MONTH12
TROUBLE_WITH_CONCENTRATION_MONTH12
DYSPHONIA_OR_APHONIA_MONTH12
COUGH_MONTH12
PEM_MONTH12
PALPITATIONS_MONTH12
PLEURITIC_CHEST_PAIN_MONTH12
SOBOE_MONTH12
BRAIN_FOG_MONTH12
CHEST_PAIN_AT_REST_MONTH12
SOB_AT_REST_MONTH12
CHEST_PAIN_ON_EXERTION_MONTH12
Dim1 (19.6%)
Dim2 (12.5%)
B Female
Dim1 (19.6%)
Dim2 (13%)
Dim1 (22.7%)
Dim2 (12%)
C Male
LOSS_OF_TASTE_MONTH3-6
LOSS_OF_SMELL_MONTH3-6
TROUBLE_WITH_CONCENTRATION_MONTH3-6
SOBOE_MONTH3-6
TROUBLE_WITH_MEMORY_MONTH3-6
MYALGIAS_ARTHRALGIAS_MONTH3-6
TROUBLE_WITH_SLEEP_MONTH3-6
HEADACHES_MONTH3-6
Dim1 (45.6%)
Dim2 (20.8%)
LOSS_OF_TASTE_MONTH12
LOSS_OF_SMELL_MONTH12
TROUBLE_WITH_CONCENTRATION_MONTH12
TROUBLE_WITH_MEMORY_MONTH12
ANXIETY_MONTH12
TROUBLE_WITH_SLEEP_MONTH12
BRAIN_FOG_MONTH12
SOBOE_MONTH12
PEM_MONTH12
SOB_AT_REST_MONTH12
WHEEZING_MONTH12
PARESTHESIAS_MONTH12
MYALGIAS_ARTHRALGIAS_MONTH12
HEADACHES_MONTH12
VISION_CHANGE_MONTH12
Dim1 (23.1%)
Dim2 (20%)

Fig.S4

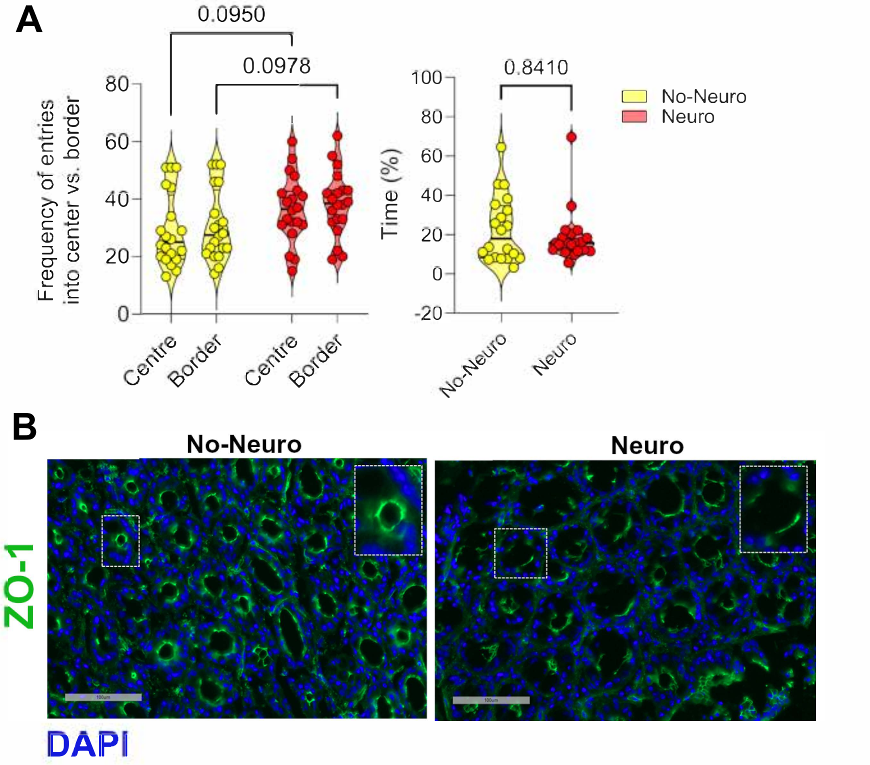
A
0.0950
0.0978
0.8410
Frequency of entries into center vs. border
80
60
40
20
0
Centre
Border
Centre
Border
Time (%)
100
80
60
40
20
0
-20
No-Neuro
Neuro
No-Neuro
Neuro
B
No-Neuro
Neuro
ZO-1
DAPI

Fig.S5

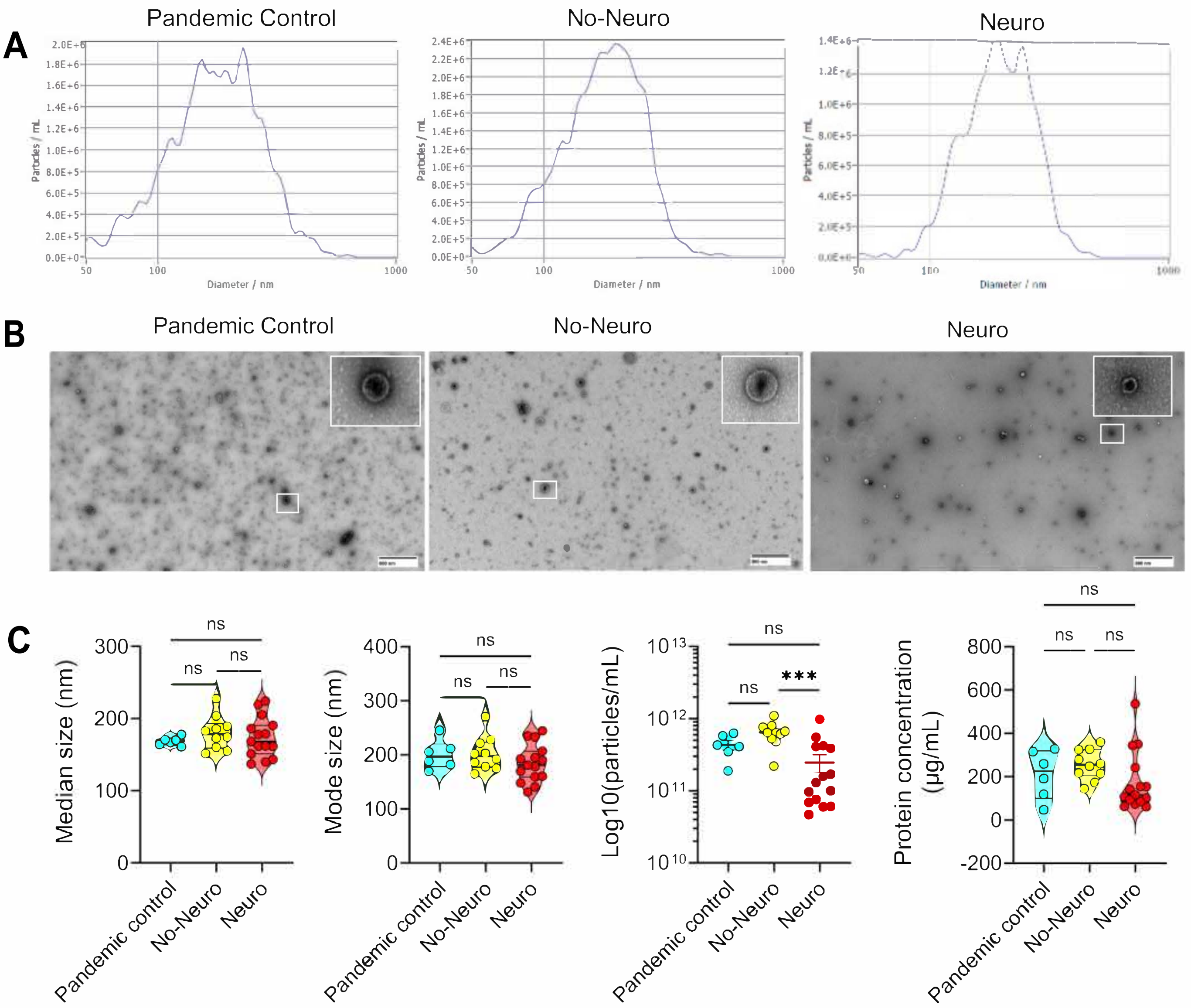
A
Pandemic Control
No-Neuro
Neuro
Particles / mL
Diameter / nm
B
Pandemic Control
No-Neuro
Neuro
C
Median size (nm)
Mode size (nm)
Log10(particles/mL)
Protein concentration (µg/mL)
ns
***
Pandemic control
No-Neuro
Neuro

Fig.S6

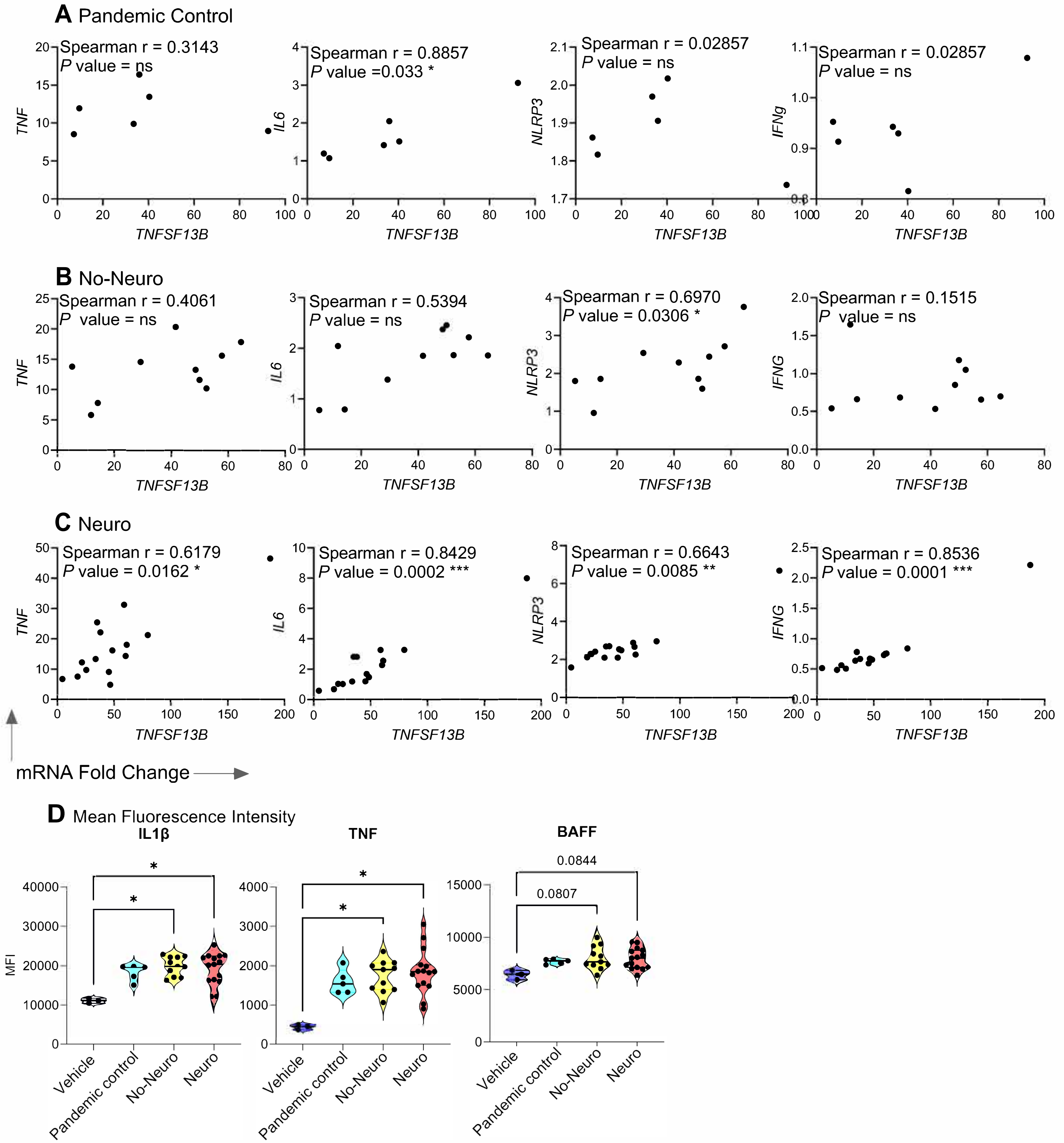

A Pandemic Control
Spearman r = 0.3143
P value = ns
Spearman r = 0.8857
P value =0.033 *
Spearman r = 0.02857
P value = ns
Spearman r = 0.02857
P value = ns
TNF
IL6
NLRP3
IFNg
TNFSF13B
B No-Neuro
Spearman r = 0.4061
P value = ns
Spearman r = 0.5394
P value = ns
Spearman r = 0.6970
P value = 0.0306 *
Spearman r = 0.1515
P value = ns
TNF
IL6
NLRP3
IFNG
TNFSF13B
C Neuro
Spearman r = 0.6179
P value = 0.0162 *
Spearman r = 0.8429
P value = 0.0002 ***
Spearman r = 0.6643
P value = 0.0085 **
Spearman r = 0.8536
P value = 0.0001 ***
TNF
IL6
NLRP3
IFNG
TNFSF13B
mRNA Fold Change
D Mean Fluorescence Intensity
IL1β
TNF
BAFF
MFI
0.0844
0.0807
Vehicle
Pandemic control
No-Neuro
Neuro

Fig.S7

A

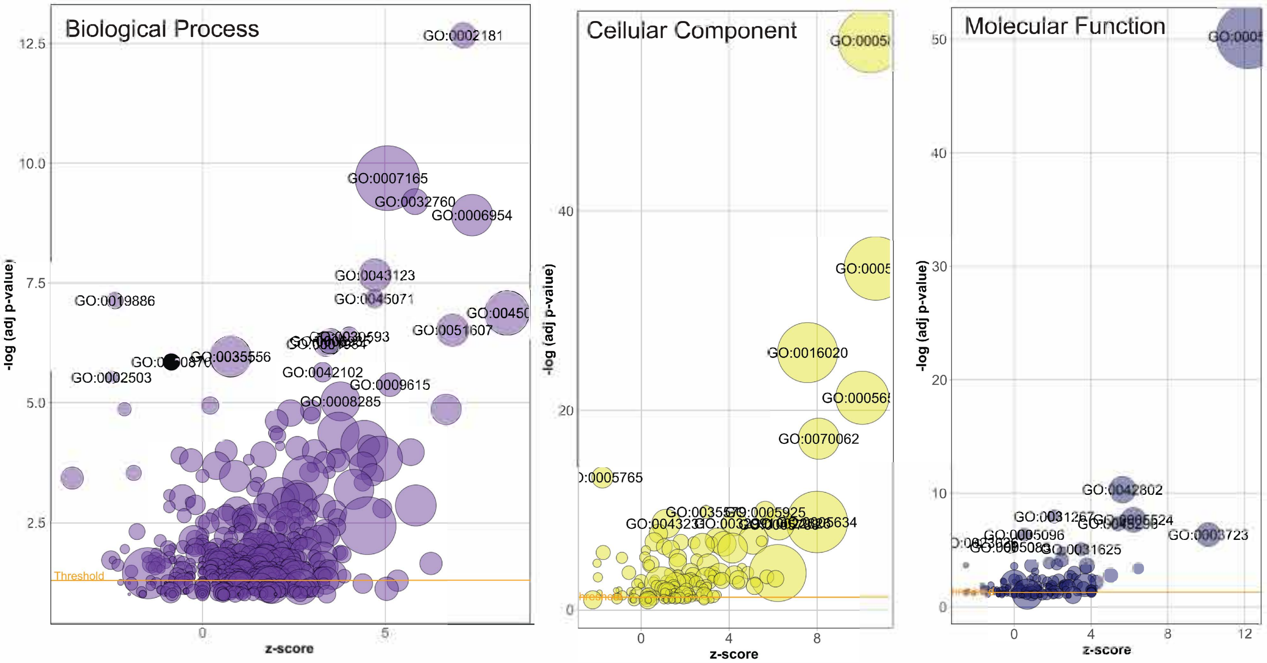


| ID | Description |
|---|---|
| GO:0002181 | cytoplasmic translation |
| GO:0007165 | signal transduction |
| GO:0032760 | positive regulation of tumor necrosis factor production |
| GO:0006954 | inflammatory response |
| GO:0043123 | positive regulation of canonical NF-kappaB signal transduction |
| GO:0045071 | negative regulation of viral genome replication |
| GO:0019886 | antigen processing and presentation of exogenous peptide antigen via MHC class II |
| GO:0045087 | innate immune response |
| GO:0051607 | defense response to virus |
| GO:0030593 | neutrophil chemotaxis |
| GO:0006935 | chemotaxis |
| GO:0001934 | positive regulation of protein phosphorylation |
| GO:0035556 | intracellular signal transduction |
| GO:0050870 | positive regulation of T cell activation |
| GO:0042102 | positive regulation of T cell proliferation |
| GO:0002503 | peptide antigen assembly with MHC class II protein complex |
| GO:0009615 | response to virus |
| GO:0008285 | negative regulation of cell population proliferation |

| ID | Description |
|---|---|
| GO:0005829 | cytosol |
| GO:0005737 | cytoplasm |
| GO:0016020 | membrane |
| GO:0005654 | nucleoplasm |
| GO:0070062 | extracellular exosome |
| GO:0005765 | lysosomal membrane |
| GO:0035579 | specific granule membrane |
| GO:0005925 | focal adhesion |
| GO:0005634 | nucleus |
| GO:0022626 | cytosolic ribosome |
| GO:0032991 | protein-containing complex |
| GO:0043231 | intracellular membrane-bounded organelle |
| GO:0005783 | endoplasmic reticulum |

| ID | Description |
|---|---|
| GO:0005515 | protein binding |
| GO:0042802 | identical protein binding |
| GO:0031267 | small GTPase binding |
| GO:0005524 | ATP binding |
| GO:0045296 | cadherin binding |
| GO:0003723 | RNA binding |
| GO:0005096 | GTPase activator activity |
| GO:0023026 | MHC class II protein complex binding |
| GO:0005085 | guanyl-nucleotide exchange factor activity |
| GO:0031625 | ubiquitin protein ligase binding |

B

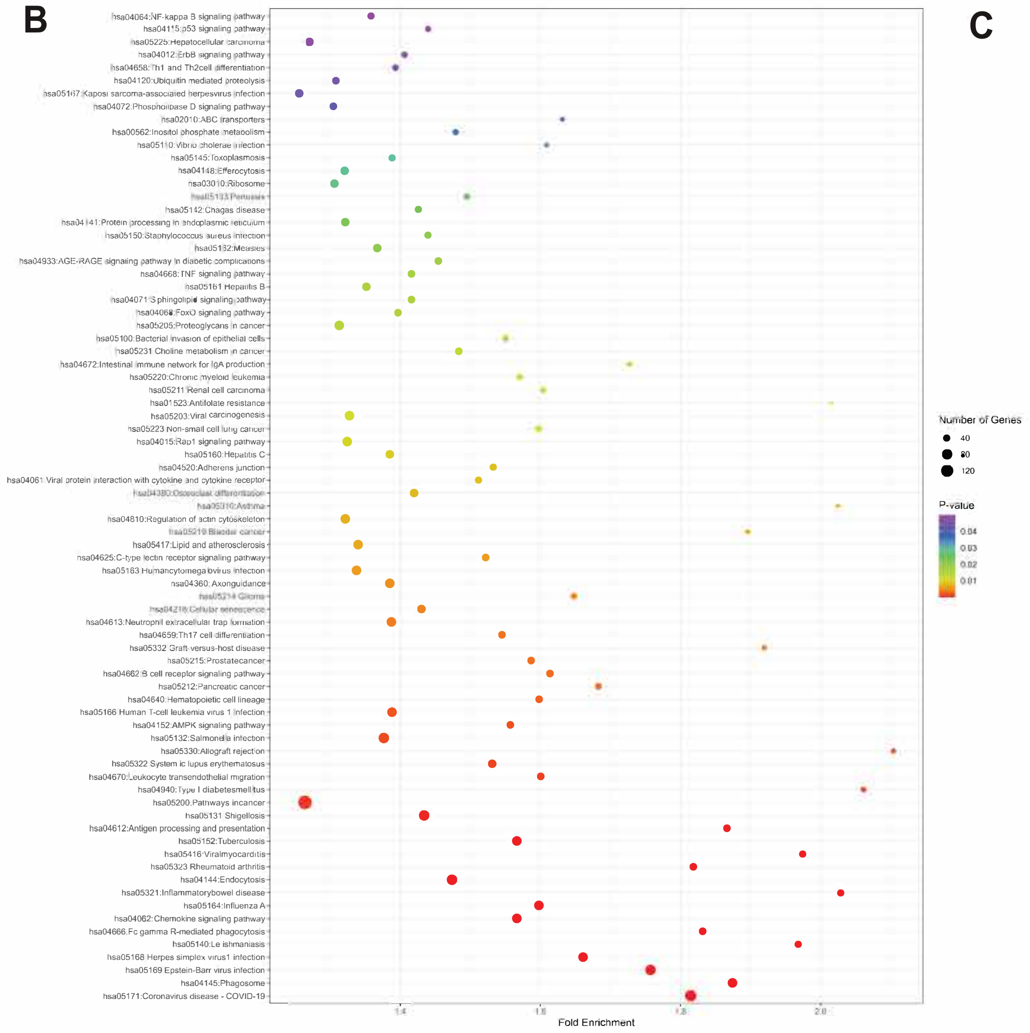


C

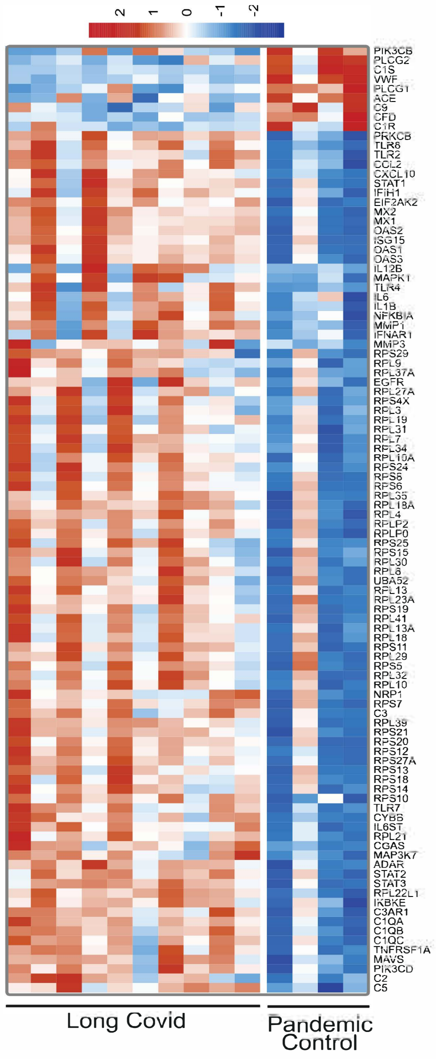

**Fig.S8**

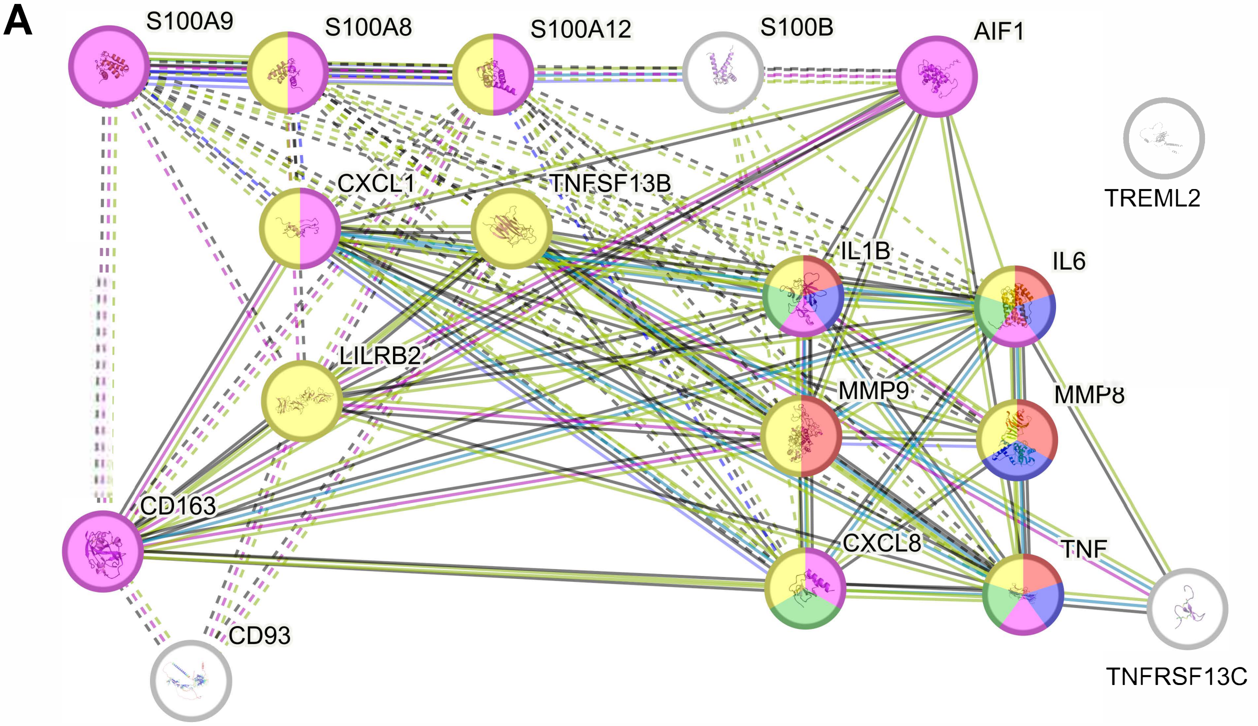


| Selected Functional Enrichments | GO-term | Description | Count in Network | Strength | Signal | False Discovery Rate |
|---|---|---|---|---|---|---|
| Biological Process | GO:0150077 | Regulation of neuroinflammatory response | 5 of 34 | 2.21 | 2.86 | 2.55E-07 |
| Biological Process | GO:0150078 | Positive regulation of neuroinflammatory response | 4 of 15 | 2.47 | 2.69 | 1.27E-06 |
| Biological Process | GO:0006954 | Inflammatory response | 10 of 538 | 1.31 | 1.85 | 4.75E-08 |
| Disease-gene-Association | DOID:0080600 | COVID-19 | 4 of 16 | 2.44 | 2.64 | 1.53E-06 |
| Tissue Expression | BTO:0001044 | Phagocyte | 11 of 135 | 1.95 | 5.23 | 1.85E-16 |

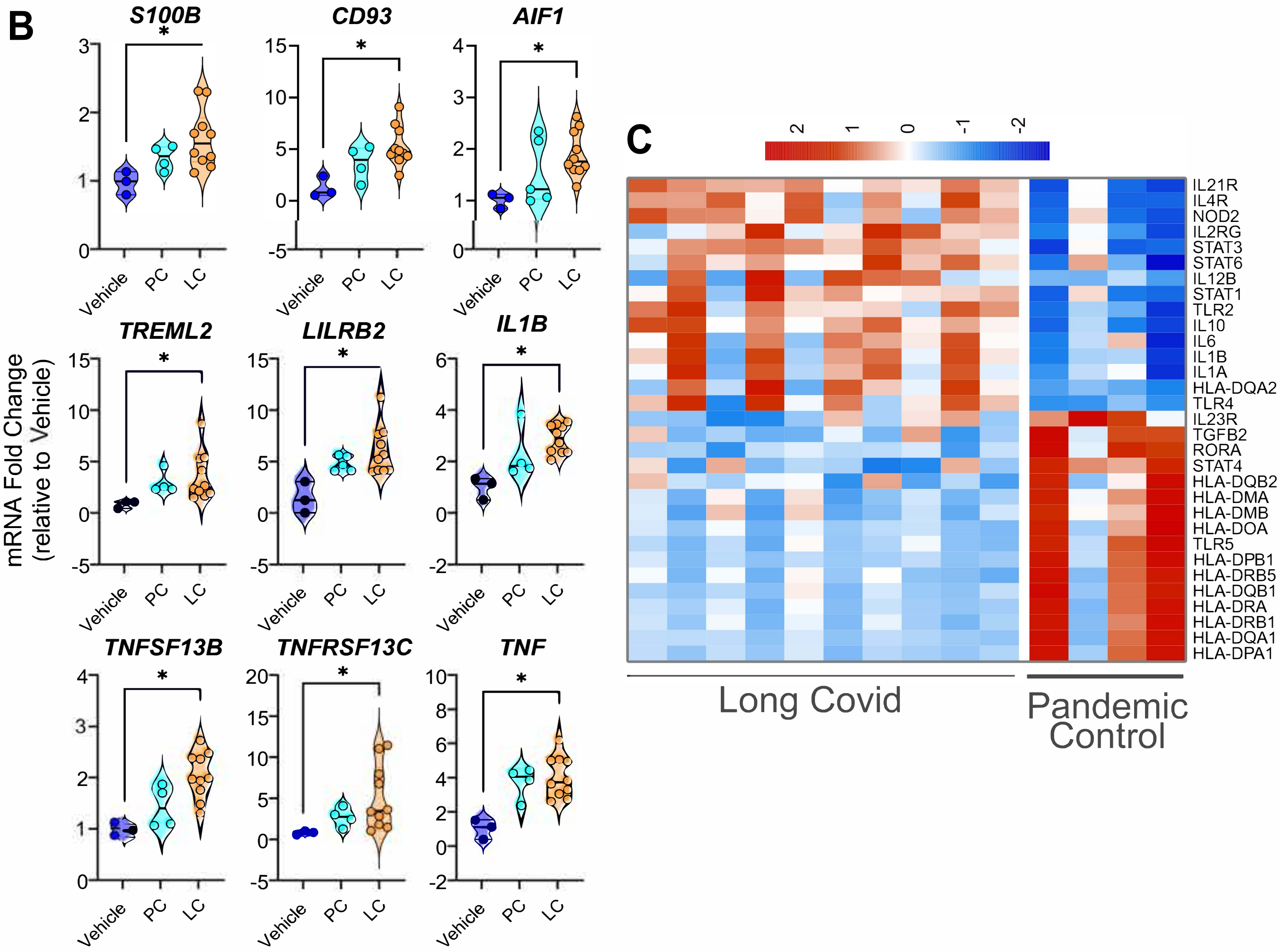

Fig.S9

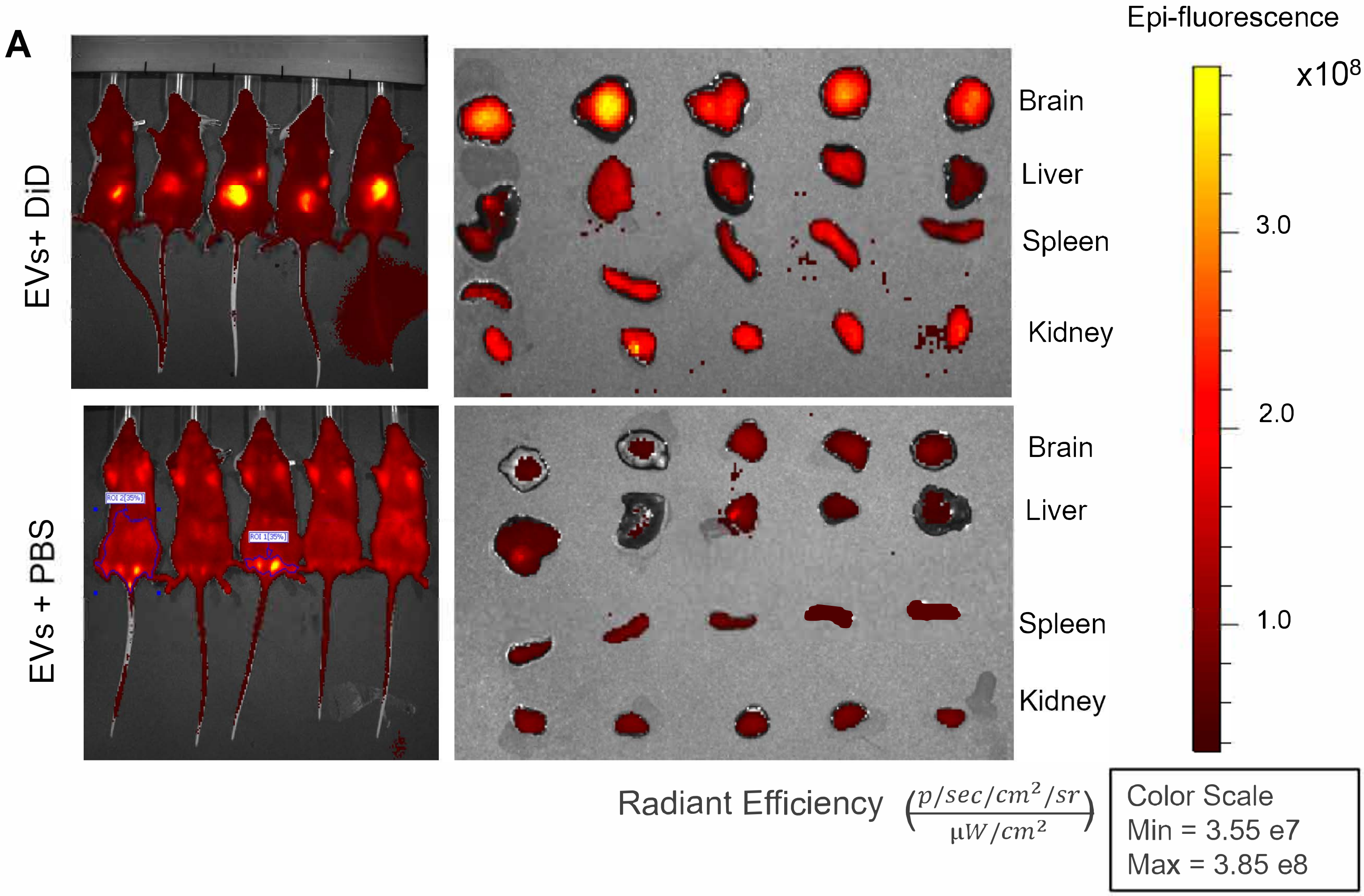
A
EVs+ DiD
EVs + PBS
Brain
Liver
Spleen
Kidney
Brain
Liver
Spleen
Kidney
Epi-fluorescence
x10^8
3.0
2.0
1.0
Radiant Efficiency (p/sec/cm²/sr / μW/cm²)
Color Scale
Min = 3.55 e7
Max = 3.85 e8

**Table S1.** Baseline characteristics of cohort participants included in microbiome analyses

| Characteristic | Pandemic controls n=12 | Long COVID n=91 | *P* value |
|---|---|---|---|
| Age, years median (IQR) | 50 (47-62) | 47 (39-58) | 0.73 |
| Female sex, n (%) | 6 (50%) | 58 (64%) | 0.54 |
| Body mass index, kg/m$^2$, median (IQR) | 29.6 (26.1-31.0) | 25.3 (22.2-28.4) | 0.034 |
| Time since SARS-CoV2 infection at enrolment, months, median (IQR) | NA | 5.4 (4.0-10.0) | NA |
| Hospitalized during acute COVID-19, n (%) | NA | 0 (0%) | NA |
| Vaccinated prior to infection, n (%) | NA | 10 (11%) | NA |
| ≥1 vaccine dose before enrolment, n (%) | 12 (100%) | 59 (65%) | 0.016 |
| Pre-existing autoimmune disease, n (%) | 0 (0%) | 16 (18%) | 0.21 |
| Pre-existing gastrointestinal disease, n (%) | 0 (0%) | 19 (21%) | 0.12 |
| Number of long COVID symptoms, median (IQR) | NA | 6 (4-9) | NA |

*P* values were calculated using Wilcoxon rank-sum test (two-sided) or Fisher's exact test, as appropriate. IQR, interquartile range; NA, not applicable.

**Table S2.** Mouse experiments and group sizes

| Experiment | Donor group (sample ID) | Neurobehavioral test | n (mice) | Sex |
|---|---|---|---|---|
| **Microbiota colonisation; experiment 1** | No-Neuro; LC1 | Open-field | 5 | Female |
| **Microbiota colonisation; experiment 1** | Neuro; LC2 | Open-field | 5 | Female |
| **Microbiota colonisation; experiment 2** | No-Neuro; LC3 | Open-field | 5 | Female |
| **Microbiota colonisation; experiment 2** | Neuro; LC4 | Open-field | 5 | Female |
| **Microbiota colonisation; experiment 3** | No-Neuro; LC5 | Open-field | 5 | Female |
| **Microbiota colonisation; experiment 3** | Neuro; LC6 | Open-field | 5 | Female |
| **Microbiota colonisation; experiment 4** | No-Neuro; LC8 | Open-field | 5 | Female |
| **Microbiota colonisation; experiment 4** | Neuro; LC7 | Open-field | 5 | Female |
| **GMEV gavage; experiment 1** | Vehicle | Y-maze/Open-field | 3 | Female |
| **GMEV gavage; experiment 1** | PC; PC1 | Y-maze/Open-field | 4 | Female |
| **GMEV gavage; experiment 1** | No-Neuro; LC3 | Y-maze/Open-field | 4 | Female |
| **GMEV gavage; experiment 1** | Neuro; LC7 | Y-maze/Open-field | 4 | Female |
| **GMEV gavage; experiment 2** | Vehicle | Y-maze/Open-field | 3 | Female |
| **GMEV gavage; experiment 2** | PC; PC2 | Y-maze/Open-field | 4 | Female |
| **GMEV gavage; experiment 2** | No-Neuro; LC1 | Y-maze/Open-field | 4 | Female |
| **GMEV gavage; experiment 2** | Neuro; LC2 | Y-maze/Open-field | 4 | Female |
| **GMEV gavage; experiment 3** | Vehicle | Y-maze/Open-field | 3 | Female |
| **GMEV gavage; experiment 3** | PC; PC3 | Y-maze/Open-field | 4 | Female |
| **GMEV gavage; experiment 3** | No-Neuro; LC8 | Y-maze/Open-field | 4 | Female |
| **GMEV gavage; experiment 3** | Neuro; LC4 | Y-maze/Open-field | 4 | Female |

All mice were female C57BL/6J. Fecal microbiota colonization experiments were performed in germ-free mice; GMEV oral-gavage experiments were performed in wild-type specific pathogen-free (SPF) mice. Vehicle, sterile phosphate-buffered saline (PBS).

**Table S3.** Primer sequences

| Primer | Species | Forward | Reverse |
|---|---|---|---|
| ***AIF1*** | **Human** | GCT ATG AGC CAA ACC AGG GA | TGG AGG GCA GAT CCT CAT CA |
| ***CD93*** | **Human** | TGA TTG CTC TCA CAG CCC AG | CAC AGA GGG GTT CCA TGA CC |
| ***CD163*** | **Human** | ACC CAG TGA GTT CAG CCT TT | AGA GAG GTG AAT TTC TGC TCC A |
| ***CXCL1*** | **Human** | TCT GGC TTA GAA CAA AGG GGC | TAA AGG TAG CCC TTG TTT CCC C |
| ***IFNG*** | **Human** | GGC TTA ATT CTC TCG GAA ACG ATG | TTC TTT TAC ATA TGG GTC CTG GCA |
| ***IL1B*** | **Human** | CCA GGG ACA GGA TAT GGA GCA | TTC AAC ACG CAG GAC AGG TAC AG |
| ***IL6*** | **Human** | CTT CGG TCC AGT TGC CTT CTC | TCA ATT CGT TCT GAA GAG GTG AGT |
| ***LILRB2*** | **Human** | TTG TCA GGG GAG CCT TGA A | CAG CCA TAT CGC CCT GTG TG |
| ***MMP8*** | **Human** | AAG CCA GGA GGG GTA GAG TT | TCC AGG TAG TCC TGA ACA GT |
| ***MMP9*** | **Human** | GGA CAA GCT CTT CGG CTT CT | TCG CTG GTA CAG GTC GAG TA |
| ***NLRP3*** | **Human** | GAT CTT CGC TGC GAT CAA CA | GGG ATT CGA AAC ACG TGC ATT A |
| ***TNF*** | **Human** | AGC CCA TGT TGT AGC AAA CC | TGA GGT ACA GGC CCT CTG AT |
| ***TNFSF13B*** | **Human** | CGG GAC TGA AAA TCT TTG AAC CAC | TGT AAG ATC CTG TTT CTT CTG GAC C |
| ***TNFRSF13C*** | **Human** | GAG AAG GGC AGG AAG GAA CC | CAA ACA CAA ACA CCC CAC CC |
| ***S100A9*** | **Human** | TCG GCT TTG ACA GAG TGC AA | GCC CCA GCT TCA CAG AGT AT |
| ***S100A8*** | **Human** | TTT CAG AAG ACC TGG TGG GG | CCC TGT AGA CGG CAT GGA AA |
| ***S100A12*** | **Human** | ATT CCT GTG CAT TGA GGG GT | GTG TCA AAA TGC CCC TTC CG |
| ***S100B*** | **Human** | ACA AGG AAG AGG ATG TCT GAG C | TGA TGA GCT CCT TCA GTT CGG |
| ***TREML2*** | **Human** | CTA CAA AAA CCG CGT GGA GG | ATC GTC CTG CAG CAA GTA GC |
| ***Gapdh*** | **Mouse** | CTG CTT CAC CAC CTT CTT GA | AAG GTC ATC CCA GAG CTG AA |
| ***Il1b*** | **Mouse** | TGA AAT GCC ACC TTT TGA CAG | CCA CAG CCA CAA TGA GTG ATA C |
| ***Il6*** | **Mouse** | TTT GCC GAG TAG ATC TCA AAG TGA | ATG CTC TCC TAA CAG ATA AGC TGG |
| ***Nlrp3*** | **Mouse** | AGA AGA GTG GAT GGG TTT GCT | GCG TTC CTG TCC TTG ATA GAG |
| ***Tnf*** | **Mouse** | CAC ACT CAC AAA CCA CCA AGT G | GAA GAG AAC CTG GGA GTA GAC AAG |
| ***Tnfsf13b*** | **Mouse** | GTC GCT CTC ATG CAA ACT GA | CCA TCA CTC CGC AGA AGG A |

**Table S4A.** Antibodies used for human macrophage flow cytometry

| Target / Host species | Fluorophore | Dilution (final) | Clone | Catalogue number | Vendor |
|---|---|---|---|---|---|
| Anti-human BAFF (mouse monoclonal) | PE | 1:10 | 121808 | IC1357P | R&D Systems |
| Anti-human IL-1β (mouse monoclonal) | APC | 1:200 | JK1B-1 | 508208 | BioLegend |
| Anti-human IL-6 (rat monoclonal) | FITC | 1:200 | MQ2-13A5 | 501104 | BioLegend |
| Anti-human TNF (mouse monoclonal) | AF700 | 1:200 | MAb11 | 557996 | BD Biosciences |

**Table S4B.** Antibodies used for immunofluorescence

| Target / Host species | Fluorophore | Dilution (final) | Clone | Catalogue number | Vendor |
|---|---|---|---|---|---|
| Anti-mouse ZO-1 (rabbit polyclonal) | N/A | 1:50 | N/A | 61-7300 | Thermo Fisher Scientific |
| Goat anti-rabbit IgG (secondary antibody) | FITC | 1:1000 | N/A | A27034 | Thermo Fisher Scientific |
| Anti-mouse GFAP (mouse monoclonal) | FITC | 1:50 | GA5 | 53-9892-82 | Thermo Fisher Scientific |
| Anti-mouse IBA-1 (rabbit polyclonal) | N/A | 1:500 | N/A | PA5-27436 | BioLegend |
